# The cow udder is a potential coinfection site for influenza A viruses

**DOI:** 10.1101/2025.08.29.673079

**Authors:** Rute Maria Pinto, Colin P Sharp, Maia Beeson, Nunticha Pankaew, Jack A. Hassard, Alexander Moxom, Callum Magill, Rebecca A. Ross, Laura Tuck, Stephen Meek, Hui Min Lee, Kirsty Jensen, Inga Dry, Pedro Melo, Jiayun Yang, Wenfang Spring Tan, Ashwin Ashok Raut, Anamika Mishra, Sjaak de Wit, J. Ross Fitzgerald, Jayne C. Hope, Joanne Stevens, Tom Burdon, Kate Sutton, Cristina L. Esteves, F. Xavier Donadeu, Ian Brown, Wendy Barclay, Thomas P. Peacock, Daniel H. Goldhill, Munir Iqbal, Pablo R. Murcia, Stuart M. Haslam, Eleanor Gaunt, Paul Digard

**Author notes:** equal contribution (Rute Maria Pinto, Colin P Sharp).

## Abstract

The incursion of high pathogenicity avian influenza A virus (IAV) into US dairy cows is unprecedented in the era of molecular diagnosis and pathogen sequencing. This raises questions over the likelihood of further outbreaks and whether dairy cattle could be a “mixing vessel” for novel strains of IAV. Using a panel of BSL2-safe reassortant viruses representing clade 2.3.4.4b H5 epizootic lineages circulating since 2020, we found that a cow B3.13 isolate displayed enhanced replication in cow mammary gland cells, along with increased viral polymerase activity and stronger interferon antagonism in cow cells compared to an earlier EA-2020-C genotype virus. However, multiple avian and mammalian IAV strains, including other clade 2.3.4.4b high pathogenicity genotypes, were replication competent in bovine cells, particularly those of the mammary gland, suggesting that there is a diverse circulating IAV pool with the potential to infect cows. Moreover, we show that cow mammary cells co-express α-2,3 and α-2,6 – linked sialic acids, and are susceptible to co-infection with human and avian IAVs. We conclude that the US cow influenza outbreak does not simply reflect a unique adaptation of the B3.13 genotype virus; rather, the bovine udder represents a permissive niche for IAV and a plausible site for reassortment, underscoring its potential role in generating novel influenza viruses with pandemic risk.

## Introduction

Since 2020, the H5N1 clade 2.3.4.4b of influenza A virus (IAV) has spread globally, threatening diverse wild bird species and causing economic devastation to the poultry industry. This panzootic has also resulted in multiple intra-phylum spill-over events, including farmed mink, semi-aquatic mammals and, in 2024, dairy cows [1, 2]). Since the detection of the B3.13 genotype H5N1 in cattle in March 2024, it has demonstrated sustained, widespread inter-state transmission in dairy cattle [3, 4]. This was unexpected, as until this event, fully verified evidence of IAV infection in cows has been limited, especially with regards to sequenced virus isolates available for further study [5]. While IAV is generally respiratory-tropic in mammals, the primary clinical indicator in the current bovine outbreak is an abrupt drop in milk production[6], correlating with high viral loads in milk[2]. Initial evolutionary and molecular analyses of cattle B3.13 viruses suggested that these viruses had quickly acquired mammalian associated adaptations in several genes [7]. Isolates were replication competent in human cell lines, were highly pathogenic in mice and ferrets [8, 9] and concomitantly, there have been multiple independent reports of zoonotic infections in dairy farm workers [10–12]. These events raised important questions around the likelihood of further IAV incursions into cattle and the pandemic potential of such outbreaks. We show here that a range of cow-derived cells are susceptible to infection with various IAV strains of avian, human or swine origin, that primary cow mammary cells express both α-2,3 and α-2,6 N-acetylneuraminic acid and that these cells can be co-infected with human and avian IAVs. We conclude that other IAV strains have the potential to infect cows and that this could yield reassortant viruses with zoonotic and pandemic potential.

## Results

### Cow mammary epithelial cells are susceptible to infection with diverse IAV strains

While IAV is traditionally considered to be a respiratory virus in mammalian species, transmission of B3.13 viruses between dairy cows has been associated with the use of milking equipment [13]. We therefore asked whether cow mammary gland epithelial cells are more susceptible to IAV infection than other cell types. Cow primary and immortalised cells originating from different anatomical sites (Figure 1A) were infected with a variety of IAV strains representative of four categories: laboratory-adapted, human, swine and avian. Infections were performed at low multiplicity of infection (MOI) and viral growth kinetic analyses were conducted up to 48 hours post infection (hpi). Of all tested cells, mammary cells showed the highest susceptibility to infection, with lab-adapted strains reaching viral titres of 10^8^ PFU/ml. Titres of 10^6-7^ PFU/ml were detected for viruses of human, swine and avian origins. Respiratory, anatomically diverse fibroblast, and immune cells were less permissive to IAV infection, with peak titres typically 2 to 3 orders of magnitude lower than those achieved in mammary cells across all virus strains (Figure 1B, Supplementary Figures S1-3). This further correlates with the ability of cow mammary cells to support the polymerase activity of diverse IAV strains (Figure 1C). The tropism of B3.13 for mammary gland epithelial cells therefore appears to be a generalisable phenomenon, observed across diverse IAV strains isolated from hosts spanning multiple taxonomic classes.

**Figure 1:**
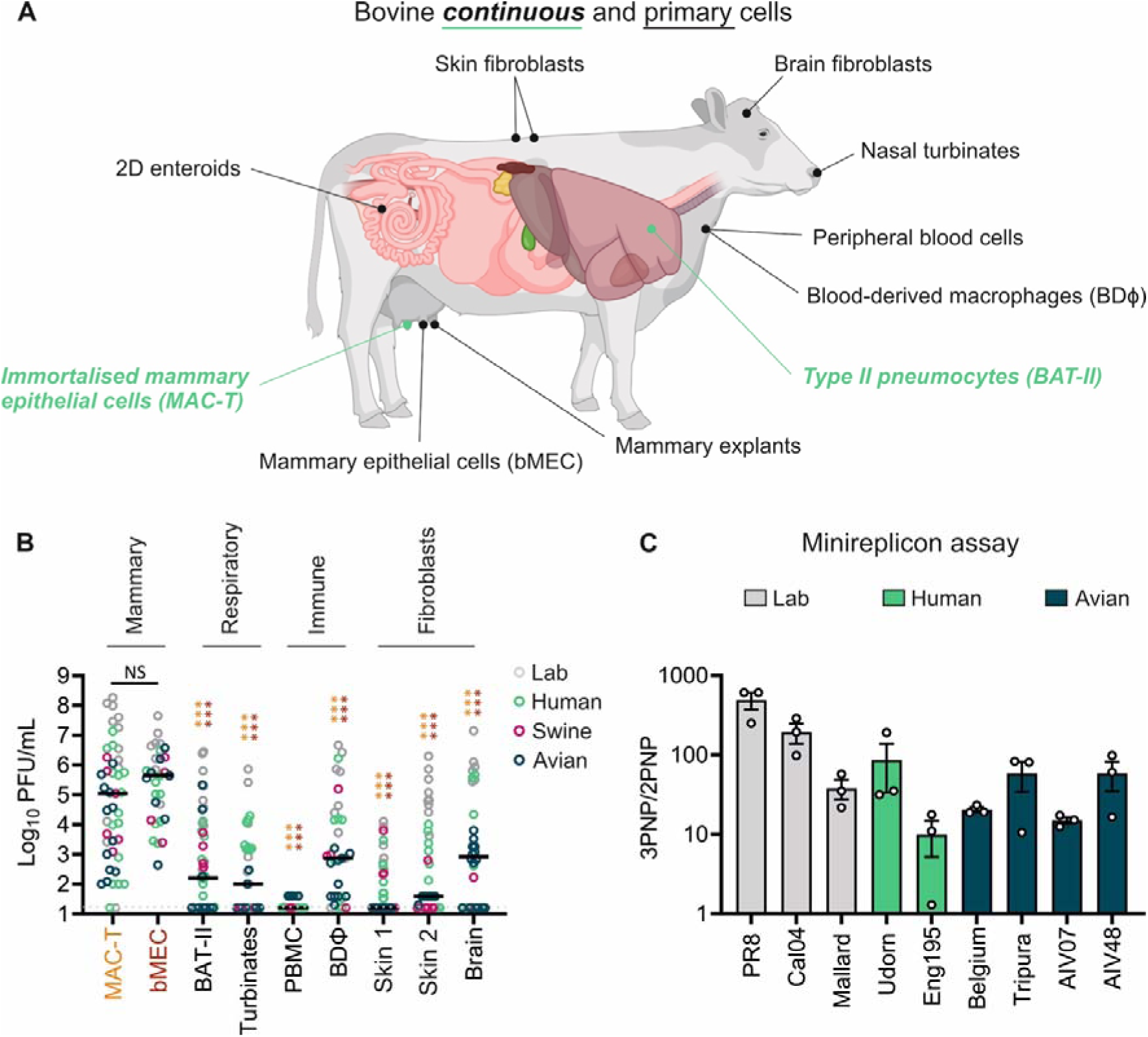
Cow cells are susceptible to infection with diverse IAV strains. A). Schematic representation of the anatomical source of immortalised (green, italic text) and primary (black) cells used in this study. B) The indicated cells were infected with a variety of viral strains at MOI 0.1 (bMEC) or MOI 0.01 (for the remaining cells) and virus titres measured by plaque assay at 48 hpi. Viruses were subcategorised into lab-adapted, human, swine and avian strains as indicated. Circles represent individual data points from 3 independently performed experiments, except PBMC (*n* = 1). Lines represent median. Statistical annotations are the result of a 2-two-way ANOVA analysis. Tukey’s multiple comparisons tests were performed against MAC-T (amber asterisks) or bMEC (burgundy asterisks) data sets. C) RNP-reconstitution assays (minireplicons) were performed in MAC-T cells by transfecting pDUAL plasmids encoding segments 1, 2, 3 and 5 from the indicated viruses alongside a firefly luciferase-expressing vRNA-like reporter plasmid. Luminescence levels were measured at 48h post transfection. Data represent mean ± SEM from three independent experiments performed in triplicate. For statistical annotations: NS denotes non significant and *** a p-value <0.001.

### B3.13 internal gene constellation confers better virus replication in cow cells

The inherent permissiveness of cow mammary cells to IAV infection does not explain why B3.13 was particularly successful at sustaining onward transmission in cows. We hypothesised that the unexpected dissemination of B3.13 infections in dairy cows could be explained by a fitness advantage of these viruses over other genotypes. To test this, biosafety level 2 viruses representative of an early B3.13 cow isolate (‘Cattle Texas’) and an avian precursor (AIV07, EURL genotype EA-2020-C) were generated harbouring the glycoproteins of the laboratory-adapted strain A/Puerto Rico/8/1934 H1N1 (PR8). Amino acid differences between the major internal gene products encoded by these viruses are summarised (Table S1). These viruses were used to infect bovine cells from different tissues including mammary epithelial cells (bMEC), type II pneumocytes, skin fibroblasts and blood-derived macrophages (BD□) as well as mammary gland explants and surface-grown intestinal organoid cultures (2D enteroids). With the exception of bMECs, where AIV07 replicated slightly better, Cattle Texas replicated to 10 to 10,000-fold higher titres compared to AIV07 (Figure 2). Therefore, Cattle Texas demonstrated a fitness advantage in bovine cells compared to a representative avian virus precursor.

**Figure 2:**
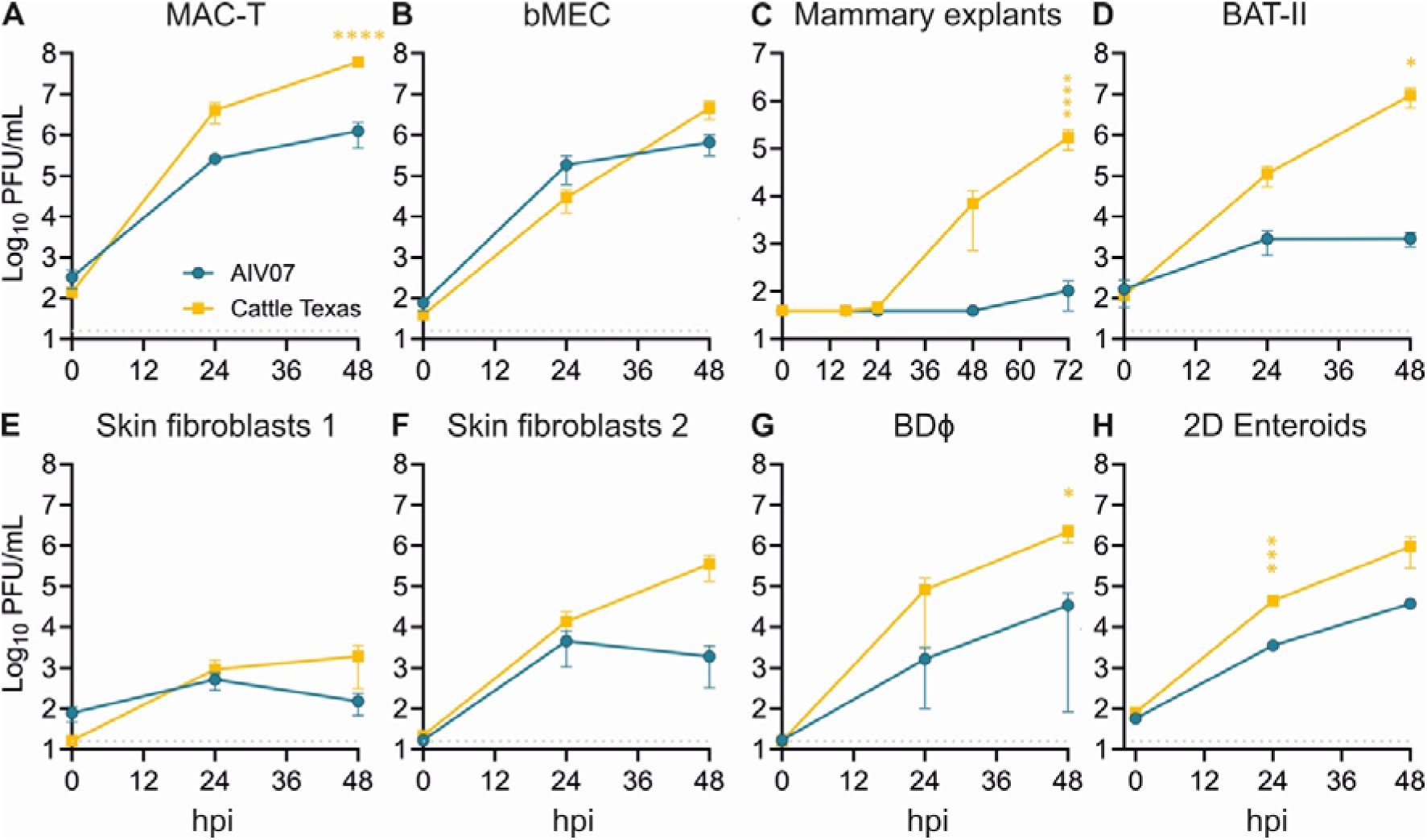
Cattle Texas is fitter in cow cells than an avian EURL genotype EA-2020-C precursor. The indicated cow cell systems were infected with 2:6 reassortant viruses with PR8 glycoproteins and internal gene constellations of either AIV07 (blue) or Cattle Texas (yellow) (MOI 0.1 in B, 5000 PFU/explant in C and MOI 0.01 for all other systems). Supernatants were collected at the indicated times post infection and virus titres determined by plaque assay. Data represent mean ± SEM from two-four independent experiments each performed in duplicate. Statistical annotations represent the results of multiple unpaired t-tests performed for each individual time post infection. Non significant results are not indicated, * indicates a p-value <0.05, *** = p-value <0.001 and **** a p-value <0.0001.Cell types were immortalised mammary epithelial (MAC-T; A), primary mammary epithelial (bMEC; B), mammary gland explant (C), immortalised pneumocytes (BAT-II; D), skin fibroblast 1 (E), skin fibroblast 2 (F), blood derived macrophage (BDΦ) (G), and 2D enteroids (H).

After entering North America through the Atlantic bird migratory route, EA-2020-C viruses like AIV07 reassorted with the local low pathogenic avian influenza (LPAI) pool, giving rise to the B3.13 genotype via acquisition of new viral genome segments 1, 2, and 5 (encoding the PB2, PB1 and NP components of the viral ribonucleoprotein (RNP) complex respectively), [14] as well as segment 8 (encoding NS1 and NS2) [7, 15, 16] (Figure 3A). Therefore, we investigated whether the new RNP segments improved viral transcription activity in cow cells using a bovine-specific minigenome reporter assay [17]. Cattle Texas RNPs were 20-times more efficient than AIV07 (Figure 3B, open blue and yellow bars). To determine which segments contributed to this improved activity, Cattle Texas and AIV07 RNPs were reconstituted with reciprocal single gene replacements, including segment 3 (encoding PA), the final component of the viral polymerase. Replacing Cattle Texas PB1 with the AIV07 counterpart had no effect, while swapping NP increased RNP activity in MAC-T cells (Figure 3B, yellow cross-hatched bars). In contrast, PB2 or PA from AIV07 significantly decreased Cattle Texas minireplicon activity. In the reciprocal swaps, AIV07 polymerase activity was improved by up to 5-fold by Cattle Texas segments 1, 2 or 3, but replacement of segment 5 had an 8-fold negative impact on polymerase efficiency (Figure 3B, green cross-hatched bars). Thus, the increased polymerase activity of Cattle Texas is attributable to the PB2- and PA-encoding segments. To test this in a viral context, we made reassortant viruses in which Cattle Texas PB2 or all polymerase segments were replaced with those of AIV07. Both PB2 and full polymerase swap viruses exhibited reduced replication in cow mammary cells (Figure 3C), confirming the importance of these segments.

**Figure 3:**
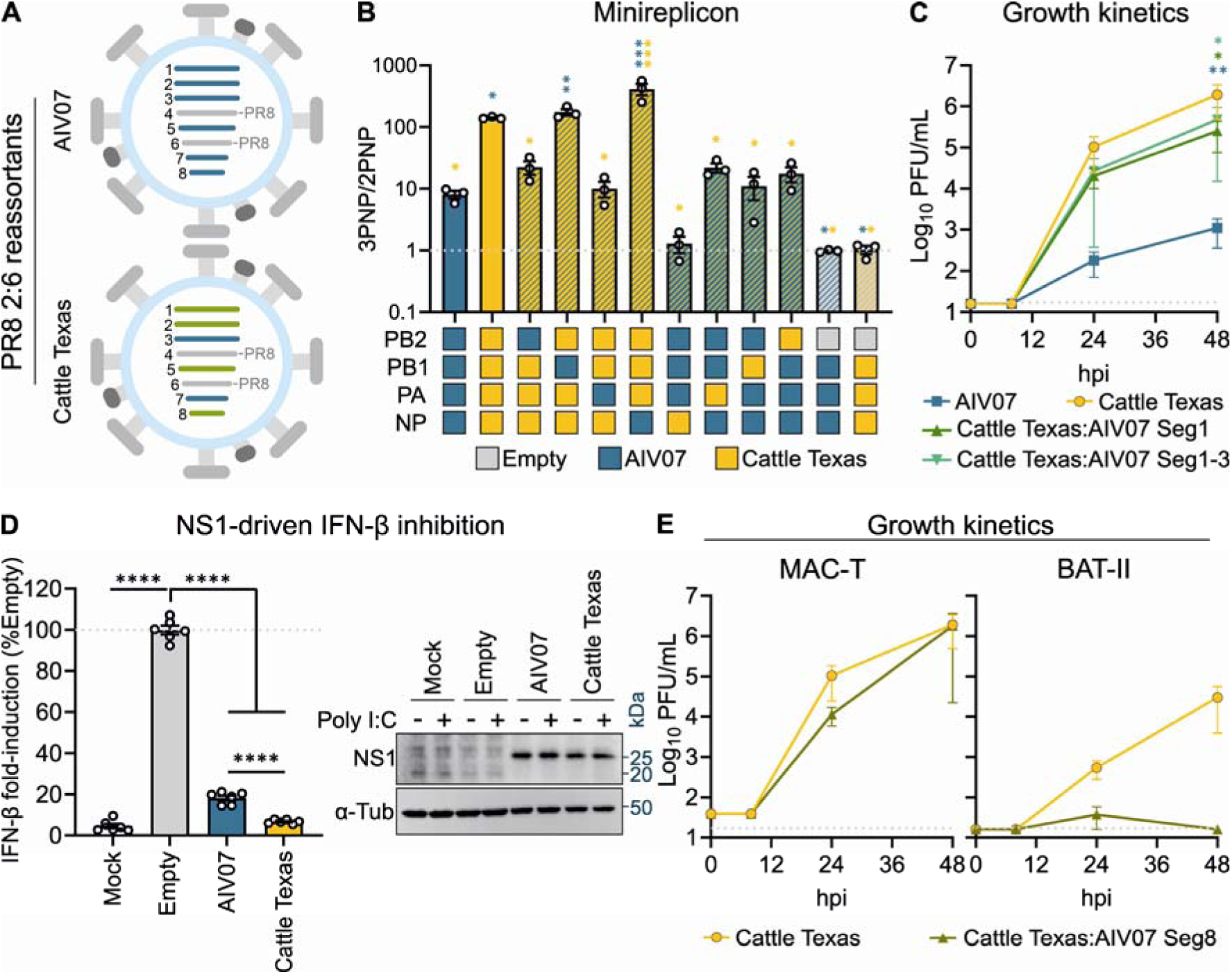
Increased fitness of Cattle Texas in cow cells correlates with enhanced viral polymerase activity and better NS1-driven inhibition of type I IFN induction. (A) Schematic representation of AIV07 and Cattle Texas segment origins. Blue and green represent European and North American ancestry, respectively. (B) RNP-reconstitution assays (minireplicons) were performed in MAC-T cells by transfecting pDUAL plasmids encoding segments 1, 2, 3 and 5 from AIV07 and Cattle Texas in the indicated combinations alongside a firefly luciferase-expressing vRNA-like reporter plasmid. Luminescence levels were measured at 48h post transfection. Data represent mean ± SEM from three independent experiments each performed in triplicate. Statistical annotations are the result of one-way ANOVA analysis. Multiple comparisons were made against full AIV07 3PNP (blue asterisks) or Cattle Texas 3PNP (yellow asterisks) using a Dunnett’s test (C) MAC-T cells were infected with polymerase-segment reassortment viruses at MOI 0.01. Supernatants were collected at the indicated time points and virus titres measured by plaque assay. Data represent mean ± SEM from three independent experiments each performed in technical duplicate. Statistical annotations are for 48h titres following a two-way ANOVA analysis and multiple comparisons using Dunnet’s test made against WT 6:2 Cattle Texas. (D) MAC-T cells were transfected with an IFN-β firefly luciferase-coding reporter alongside AIV07 or Cattle Texas segment 8 plasmids, and poly I:C treated 24h later. Luminescence readings were acquired after a further 24h. Data (left hand panel) represent mean ± SEM from three independent experiments each performed in technical triplicate. Statistical annotations are the result of one-way ANOVA analysis followed by a multiple comparison Dunnett’s test. A western blot (right hand panel) was also performed to detect NS1, using α-Tubulin as a loading control. The positions of molecular weight markers are indicated. (E) MAC-T cells were infected with segment 8 reassortant viruses at MOI 0.01. Supernatants were collected at the indicated time points and virus titres measured by plaque assay. For all shown statistical annotations: NS indicates non significant, * a p-value <0.05, ** a p-value <0.01, *** a p-value <0.001 and **** a p-value <0.0001.

NS1 is the primary IAV IFN antagonist, working through multiple partially redundant but also strain-specific mechanisms, including both general repression of cellular gene expression at transcriptional and translational levels as well as specific inhibition of IFN induction [18]. As an initial test of whether reassortment of segment 8 was phenotypically important, we assessed host cell shut off by infecting MAC-T cells with the PR8 6:2 Cattle Texas or a PR8 2:5:1 Cattle Texas harbouring AIV07 segment 8 and pulsed the cells with puromycin at various times post infection. Western blotting for the puromycin-labelled prematurely terminated polypeptides indicated that while both viruses repressed cellular translation at late times post infection, there was no major difference between them (Figure S4A). As a more quantitative measure of shut off, we co-transfected MAC-T cells with segment 8 plasmids from the two viruses alongside a constitutive expression plasmid encoding *Renilla* luciferase and measured the resulting luminescence. Both Cattle Texas and AIV07 segment 8 gave dose-dependent inhibition in this assay with Cattle Texas showing a modest improvement over AIV07 (Figure S4B). The PA-X accessory gene expressed from segment 3 also contributes to IAV host cell shut off [14]. Although segment 3 reassortment did not occur between genotypes EA-2020-C and B3.13, genetic drift led to four amino acid differences between the PA-Xs of AIV07 and Cattle Texas, three of which lie within the “X-ORF” (Figure S4C). Therefore, we similarly compared the ability of AIV07 and Cattle Texas PA-X proteins to inhibit host gene expression, but no significant differences were observed (Figure S4D).

To test specific IFN antagonism by NS1, the AIV07 and Cattle Texas segment 8 plasmids were transfected into MAC-T cells alongside an IFN-β reporter plasmid and the cells stimulated with poly I:C as a mimic for viral RNA. This showed that, at similar expression levels, Cattle Texas segment 8 had a significantly higher ability to counteract type I IFN induction than that of AIV07 (Figure 3D). To assess the contribution of this to overall virus fitness, replication of the 2:5:1 Cattle Texas reassortant virus harbouring AIV07 segment 8 was characterised in MAC-T and BAT-II cells. Introduction of AIV07 NS into Cattle Texas resulted in a slight but non-statistically significant delay in virus propagation in mammary epithelial cells, but had a much larger effect in the pneumocyte cell line (Figure 3E), suggesting that changes in segment 8 also contribute to the improved fitness of Cattle Texas over AIV07. Thus overall, the reassortment events that replaced segments 1 and 8 alongside genetic adaptations in segment 3 may have contributed to the ability of the B3.13 genotype virus to infect cattle.

Spill-back of the cattle B3.13 virus into poultry has been observed on multiple occasions [3, 19], despite its evident fitness adaptations for bovines. The 2:6 Cattle Texas virus replicated to slightly lower titres in chicken lung epithelial cells than AIV07, while its polymerase and NS1/NS2 exhibited similar activities to those of the AIV07 proteins in the same cells (Supplementary Figure S5). Thus, the enhanced fitness of Cattle Texas in bovine cells is specific to bovine cells but does not carry a high fitness penalty in chicken cells.

### Cow mammary cells are susceptible to European 2.3.4.4b genotype viruses and support co-infection by human and avian viruses

Detections of IAV in US dairy cattle are mostly of North American H5N1 2.3.4.4b genotypes B3.13 and, more recently, D1.1 viruses. Therefore, we asked whether similar dairy cattle incursions could happen in countries where the B3.13 and D1.1 genotypes are not circulating. To that end, we compared the growth kinetics of Cattle Texas with other avian European 2.3.4.4b genotypes in primary and continuous bovine mammary cells and explants, again all as 2:6 reassortants with PR8 glycoproteins to permit work at BSL2. In addition to AIV07 (genotype EA-2020-C), these included a 2.3.4.4b 2020 H5N8 precursor (H5N8-20) and EURL genotypes EA-2021-AB (AIV09) and EA-2022-BB (AIV48) (Figure 4A). In all three systems, Cattle Texas replicated to the highest titres, perhaps as expected for a virus with bovine-adapted internal segments. Other unadapted 2.3.4.4b strains showed varied viral fitness (Figure 4A). Cow mammary explants showed the greatest range of virus titres, with H5N8-20 and AIV07 exhibiting very poor fitness in this model, such that many of the explants were refractory to infection or only yielded low titres (Figure S6A-D). However, the more recent AIV09 and AIV48 isolates replicated better in the explants, and Cattle Texas demonstrated the highest virus fitness, consistent with a trend for increasing replication fitness in mammary tissue of the 2.3.4.4b strain over time. This suggests that recent Clade 2.3.4.4b H5N1 genotypes on both sides of the North Atlantic have the potential to cause dairy cattle infections.

**Figure 4:**
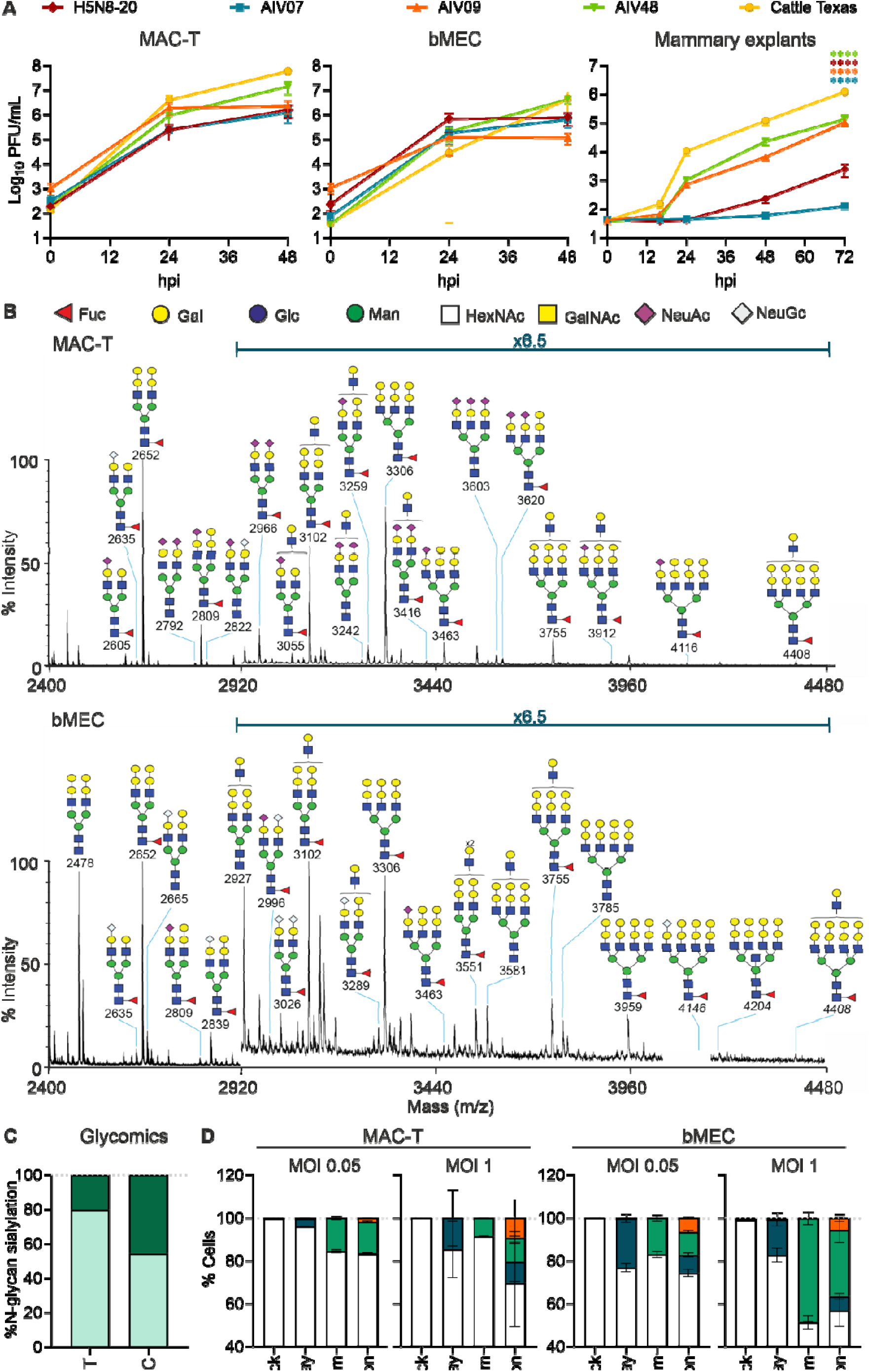
Cow mammary cells are infectable with viruses of human and avian origin. (A) MAC-T (MOI 0.01), bMEC (MOI 0.1) cells and mammary explants (5000 PFU/explant) were infected with the indicated PR8 2:6 viruses. Supernatants were collected at the indicated times post infection and virus titres determined by plaque assay. Statistical analysis was performed by a 2-way ANOVA and multiple comparisons were performed against Cattle Texas in specific time points using a Dunnett’s test. (B) Annotated MALDI-TOF spectra of permethylated N-glycans from MAC-T and bMEC cells. Annotations show [M + Na]^+^ molecular ions. Peak annotation is based on composition, biosynthetic knowledge and MS/MS analysis. (C) Proportions of a-2,3 and a-2,6 linked sialic acid in MAC-T and bMEC N-glycans. (D) MAC-T and bMECs were infected with combinations of Norway (pH1N1) and/or Belgium (H3N1) at MOI 0.05 for 48h or MOI 1 for 16h. Infections at MOI 0.05 were overlayed with media containing TPCK-treated trypsin. Cells were fixed, stained with anti-H1 and H3 before quantification was performed by FACS analysis. Data represent mean ± SEM from three independent experiments each performed in technical duplicate. For all shown statistical annotations: ****p-value <0.0001.

Human IAV pandemics classically arise from reassortment between avian and mammalian strains [20, 21]. Historical data suggest the incursion of human seasonal strains into cattle [5], so in the face of an ongoing H5N1 epidemic in birds and cattle, the possibility of the cow udder as a “mixing vessel” arises. However, co-infection of a single cell harbouring both avian (α-2,3 linked sialic acid) and mammalian (α-2,6 linked sialic acid) receptor types is needed for this mixing to occur. Lectin staining and mass spectrometry analyses indicate the presence of both α-2,3 and α-2,6-linked sialic acid receptors in cow mammary glands [22–25]. To establish the amounts of each receptor type in the cells used here, we next performed quantitative glycomic analyses on the immortalised and therefore likely relatively homogeneous (MAC-T) and primary, potentially heterogeneous (bMEC) cell populations. This identified complex N-glycans containing a mix of α-2,3 and α-2,6 linked sialic acids, with a higher proportion of α-2,6 linkages present on the bMECs compared to MAC-T cells (Figure 4B, C; Supplementary Figures S7-S10). Cow mammary cells therefore express the sialic acids required for infection with both avian and mammalian IAVs; however, this does not demonstrate that the two sialic acid conformations are present at functionally relevant levels on the same cell. To determine whether individual cow mammary gland cells could be co-infected by human and avian strains of virus, MAC-T and bMEC cells were inoculated with an α-2,6 sialic acid-dependent human seasonal H1N1 strain and an α-2,3 sialic acid-dependent low pathogenicity avian virus (both of which replicate in mammary explants (Figure S6E-H), the most discriminatory bovine cell system for replication of the 6:2 reassortant HPAIVs (Fig 4A)), either singly or together and at low or moderate MOI. Infected cells were fixed and stained with anti-H1 or H3 HA and analysed by flow cytometry. The individual infections confirmed antibody specificity, while dual-infected cell populations were clearly identified in all co-infection samples (Figure 4D). We therefore conclude that co-infections of human and avian viruses within the cow mammary glands are possible, supporting the possibility of reassortment events.

## Discussion

The unexpected influenza outbreak in US dairy cows requires careful consideration to understand why it occurred, and its potential implications for both livestock and human health [15]. At one extreme, it could have resulted from a rare adaptation of the B3.13 genotype to allow infection of a species usually refractory to infection with most strains of IAV; at the other, an unusually large and well documented epidemic of a not uncommon spillover event that mostly goes undetected. To experimentally test these possibilities, we compiled a variety of bovine cell culture systems and performed a systematic analysis of their susceptibility to IAV infection. We found that while there are clear virus strain and cell-type differences in the susceptibility of bovine cells to IAV infection, many cell types are susceptible to infection with a diverse range of avian and mammal viruses. Notably however, we think our data support mammary gland epithelial cells to be the primary site of infection compared to respiratory, immune or fibroblast cells, consistent with the clinical manifestations of mastitis and high viral loads present in milk [4, 26]. However, the lack of a wide selection of epithelial cell types from other anatomical sites in the cow means that although we can be confident that bovine mammary cells are highly permissive to IAV infection, we cannot rule out that other organs might be equally or even more susceptible if epidemic transmission were to occur by another route than milking equipment. The bovine cells used in our study come from a mixture of dairy and beef cattle. The mammary explants were derived from dairy cattle of unspecified breeds, while MAC-T cells are from Holstein Friesian dairy cattle. The bMECs were derived from beef cattle, and the enteroids from male Holstein Friesian calves. We can therefore be confident that *in vitro* cell permissivity to IAV infection is not unique to a single breed, sex or age of cattle, but we cannot rule out that these factors might influence viral susceptibility *in vivo* [27].

The ability of cow mammary cells to be infected with and support high levels of replication of a wide range of IAV strains here described argues against the extreme hypothesis of a truly unique phenotypic trait of the B3.13 strain that permits infection of cows. These observations are consistent with recent studies showing that other avian 2.3.4.4b isolates are able to replicate in bovine cells [28–30] and udders [26, 31], as well as the more recently detected D1.1 genotype spillover into US dairy cattle [32]. However, our findings of wide variation in the ability of specific strains of IAV to replicate in bovine cells suggests that adaptation does play a role. Thus, we sought to determine what, if anything, predicted the emergence and transmission of Cattle Texas above other avian IAV strains. We found a multifactorial scenario, with improvements in polymerase efficiency and the ability of segment 8-encoded products (most likely NS1) to counteract type I interferon induction. These improvements were apparent in cow but not chicken cells, indicative of evolutionary adaptations to the new host. This is consistent with other studies showing a replicative advantage of cattle isolates over an avian precursor [8, 31]. For PB2 and PA, the amino acid mutations PB2 M631L and PA K497R present in the Cattle Texas strain we used have recently been reported to enhance polymerase efficiency and virus replication in cow mammary cells [3, 17, 19]. Furthermore, impairment to transcriptional efficiencies rendered by the newly acquired segment 5 imply that there is still the potential for further fitness gains of the B3.13 genotype in cattle. Notably, the D1.1 strain isolated from cows in Nevada possess PB2 D701N [33] while a human isolate contained PB2 E627K [34], suggesting that other adaptation pathways in cattle are possible [35, 36].

Pigs are the classic reassortment host of IAV, the respiratory tract of which is unusual in its ability to support infection with both avian and mammalian IAV strains and from which the most recent influenza pandemic in 2009 arose [37, 38]. However, HPAI H5 infections in pigs are rare; pigs were either not susceptible to H5N1 2.3.4.4b strains following nasal exposure [39], or when infected no transmission was observed [40]. Therefore, reassortment of H5 HPAI with other mammalian IAV strains within the swine respiratory tract can be considered unlikely. However, our observations on the ability of avian and mammalian IAV strains to infect cow mammary glands raised the question of whether cow udders could serve as tissues permissive for viral reassortment for the generation of novel strains containing human-adapted and H5 HPAI-origin segments, analogous to the pig reassortment events that are conjectured to have precipitated previous pandemics. The fact that primary cow mammary cells can be co-infected with human and avian IAV strains, supported by the mixed distribution of α-2,3 and α-2,6 linked sialic acid observed in these cells, suggests that cow udders do have this potential. However, our experiments necessitated culture of these cells and culture passage has been observed to increase overall sialylation in some cell types [41, 42], meaning that our results may modestly overestimate the total sialylation abundance.

Infection of cows with human and other IAV strains has been suggested in the past. In 1997, dairy cows in the southwest of England presented with sporadic drops in milk production, which the reporting authors considered to be attributable to IAV [43, 44]. Additional studies have reported seroconversion against circulating seasonal H3N2 virus in British cattle, which was associated with reduced milk yield and respiratory disease [45, 46]. Despite the occasional respiratory signs, such as nasal discharge and increased respiratory rate followed by seroconversion, virus isolation from respiratory swabs from these seroconverted animals was unsuccessful. With hindsight, isolation from milk may have had a higher chance of success. Consequently, the idea that “cows could foster flu pandemics” was considered back in the early 2000s [47]. Moreover, the recent seroconversion for H5N1 2.3.4.4b in sheep and detections of IAV RNA in milk [48, 49] further suggests that other ruminant livestock species may hold similar potential. Until recently, the absence of H5 detection in cattle beyond the United States might have indicated that the coordinated polymerase and NS1 and/or NS2 adaptations that we have linked to this type of spillover have not yet emerged in Europe or other regions. It could also be attributable to insufficient surveillance, potentially due to the unexpected presentation as mastitis. However, H5 seropositive dairy cattle in the Netherlands have recently been reported [50]. At the time of writing, no viral sequences have been made available, so whether this represents similar viral adaptation to what we describe for the B3.13 genotype and/or enhanced surveillance remains to be determined.

Overall, our findings point to requirements for reconfigured and enhanced policy procedures and surveillance of IAV incursions into ruminant livestock species. Currently, mastitis outbreaks are not routinely investigated for the presence of IAV. Adding IAV testing to mastitis diagnostic assessments as a differential when other common causes have been excluded will be beneficial to livestock. This will allow early detection of emerging events with implementation of appropriate infection control measures and justify contact tracing of farm workers.

## Materials and Methods

### Ethics

All reverse genetics work used a loss-of-function approach and was carried out under a license (GMRA1811) from the UK Health & Safety Executive. All work with avian viruses was carried out in a segregated BSL2 laboratory in the Roslin Institute, separate from where mammalian IAV strains are handled.

All animal experimental work was completed under license and in accordance with the Animal (Scientific Procedures) Act 1986 and with approval from the Roslin Institute animal welfare and ethics committee under the project licence number PP0612543. All studies were performed in line with the Animal Research: Reporting of *In Vivo* Experiments (ARRIVE) Guidelines [51].

### Cell line maintenance, primary cell isolation, derivation and culture

Madin-Darby Canine Kidney (MDCK) (American Type Culture Collection, ATCC), MDCK expressing galline ANP32A (MDCK-GgANP32A) [52], MDCK expressing Sialyltransferase 1 (MDCK-SIAT) [53], human embryonic kidney (HEK293T), immortalised mammary epithelial cells (MAC-T) [54] and bovine nasal turbinate (BT) cells (ATCC, CRL-1390; predominantly fibroblast morphology) were cultured in DMEM supplemented with 10% foetal bovine serum (FBS), 2mM glutamine, 100 U/ml penicillin, and 100 μg/ml streptomycin. MDCK-SIAT cells were further supplemented with 50 µg/ml geneticin. MAC-T cells were further supplemented with 5 µg/ml insulin and 1 µg/ml hydrocortisone. Chicken lung cells (CLEC213) [55] were cultured with DMEM/Nutrient Mixture F-12 also supplemented with 8% FBS, 100 U/ml penicillin and 100μg/ml streptomycin. BAT-II cells [56, 57] were maintained in Small Airway Growth medium (Lonza) further supplemented with 10% FBS, 100 U/ml penicillin and 100μg/ml streptomycin. All immortalised cells were regularly tested for mycoplasma contamination and passaged once or twice weekly.

Primary cow mammary epithelial cell (bMEC) lines were generated from the mammary glands of Aberdeen Angus cross cows as previously described [58]. Glands were sprayed with 70% ethanol and skin was removed. Cuboidal epithelial tissue fragments (5 cm in length) were cut from the gland, submerged in 70% ethanol for 30s and washed 5 times in PBS containing antibiotics and 2.5 μg/ml amphotericin B. Tissue fragments were transferred to a glass petri dish and minced with scalpels to 2-3 mm pieces. Next, 10-15g of minced tissue was dissociated overnight with low shaking in 20 ml of DMEM/Nutrient Mixture F-12 supplemented with 15 mM HEPES and 0.075% sodium bicarbonate, 2% Bovine serum albumin (BSA), 300 U/ml Collagenase Type I and 100 U/ml Hyaluronidase from bovine testes. Dissociation media was removed by centrifugation and pellet was washed twice with DMEM/F12. This yielded small aggregates of part-digested tissue which were resuspended in freezing medium and stored at −150°C. To obtain a single cell suspension, aliquots of part-digested tissue aggregated were thawed, washed with Hanks’ Balanced Salt Solution (HBSS) supplemented with 2% FBS, resuspended in warm trypsin and vigorously pipetted for 3 mins and washed with HBSS/2% FBS, resuspended in a warm Dispase/DNAse solution and gently pipetted for 3 minutes. Ice cold HBSS/ 2% FBS was added and the suspension was filtered on a 70 μM strainer, cells were washed by centrifugation, resuspended in growth media (DMEM/F12 supplemented with 0.1% BSA, 10 ng/ml epidermal growth factor (EGF), 10 ng/ml cholera toxin, 1 µg/ml bovine insulin, 0.5 µg/ml hydrocortisone, 100 U/ml penicillin, 100 μg/ml streptomycin and 250 µg/ml gentamycin) and seeded onto collagen coated wells/flasks. bMECs were examined everyday by light microscopy to check for signs of contamination, viability, morphology, and confluence. Media was refreshed daily. Growth media was further supplemented with 2.5 µg/ml amphotericin B, the concentration of which was halved every day until the cells had reached sufficient confluency to be expanded. Initial tests of the permissivity of bMECs to infection with IAV (PR8 and the Cattle Texas 6:2 reassortant) indicated that 0.01 MOI infections gave more variable outcomes than infections at an MOI of 0.1 (Figure S11), perhaps because of heterogeneity in the cell populations. The average end point (48h) titres were not significantly different between the two MOIs, so for further experiments an MOI of 0.1 was used to increase the reproducibility of the data.

Primary fibroblasts were extracted from skin (two independent preparations; referred to as ‘skin 1’ and ‘skin 2’) and brain tissue explants dissociated with 5 mg/ml Collagenase type I at 37°C, shaking for 3h. The dissociated material was passed through a 70 µm filter and washed with HBSS. Cells were cultured on bovine gelatine-coated tissue culture plastic and expanded in Glasgow’s MEM) supplemented with 10% foetal bovine serum, 2 mM L-glutamine, 1x MEM non-essential amino acids, 1 mM sodium pyruvate, 0.1 mM 2-mercaptoethanol, 100 U/ml penicillin-streptomycin, 50 µg/ml gentamicin and 2.5 µg/ml amphotericin B. Established cultures were maintained in the same medium without antibiotics.

Cow PBMCs were collected from Holstein-Friesian cattle housed at the University of Edinburgh farm using published procedures [59]. Cow blood derived macrophages (BDΦ) were generated from PBMCs as described previously [59] except that the macrophages were cultured in BDΦ medium (RPMI-1640 supplemented with 20% FBS, 4 mM L-glutamine and 50 µM β-mercaptoethanol, 100 U/ml penicillin-streptomycin and 20 ng/ml His-tagged recombinant bovine CSF1 - the latter supplied by the Roslin Immunological Toolbox). BDΦ medium was replaced on day 7. On day 12 the adherent cells were rigorously washed with phosphate-buffered saline PBS and detached with TrypLE Express. Purified BDΦs were resuspended at 1.5 x 10^5^ cells/ml in BDΦ medium without CSF1, aliquoted into 24 well plates (1ml/well) and cultured for 48h before infection on day 14.

Bovine 3D enteroids were derived from five 6-month-old healthy male British Holstein–Friesian calves housed at the University of Edinburgh farm. Calves were subject to Schedule 1 method of cull and sections of distal ileum were obtained post-mortem and stored in HBSS on ice until further use. Crypts were isolated and 3D enteroids were cultured as previously described [60]. Bovine 3D organoid cultures were used to generate 2D enteroids as previously described [61]. The 2D enteroids are comprised of predominantly epithelial cells including Paneth, goblet and enteroendocrine cells, as well as stem cells.

Bovine mammary gland explants were extracted from udder tissue collected at Sandyford Abattoir (Paisley, Scotland) from healthy cows of unspecified breeds that had been culled for the food chain. Excess tissue was removed, teat and gland cistern were dissected (Figure S12A) and transferred to transport media comprising of chilled DMEM supplemented with 100 μg/ml Penicillin/ 100 μg/ml streptomycin and 5 μg/ml fungizone. Every 45 minutes, tissues were transferred to 500 ml of fresh transport media. Inside class II biological safety cabinet, explants were made using a 5 mm biopsy punch. Biopsies were added to 24 well plates with 500ul of transport media further supplemented with 10% FBS and incubated for 24h at 37°C, 5% CO_2_ prior to infection. Histological examination of the explants showed that they consist of duct epithelium over successive layers of connective and muscle tissue (Figure S12B).

Validation of bovine origin of the primary and continuous cell lines was performed by PCR. DNA was extracted from cell pellets, and the cytochrome B gene was amplified by PCR with degenerate primers (CGAGACGTVAAYTAYGGMTGA and ACBGTDGCYCCTCARAADGATATTTG). Cycling conditions were 94°C for 2 minutes, followed by 50 cycles of 94°C for 18 seconds, 50°C for 21 seconds then 72°C for 30 seconds with a final extension of 72°C for 5 minutes. Amplicons were sent for Sanger sequencing and resultant sequences were aligned to the Bos taurus mitochondrion complete genome reference sequence (NCBI accession number NC_006853.1).

All cells, organoids and explants were kept at 37°C and 5% CO_2_ under normoxic conditions.

### Viruses

Selected viruses were split into 4 different categories based on their history and isolation animal host (Table 1). A/Puerto Rico/8/1934 H1N1 (“PR8”) [62], mouse adapted A/California/04/2009 H1N1 (“Cal04”) [63], A/WSN/33 H1N1 (“WSN”), swine adapted A/mallard/Netherlands/10-Cam/1999 H1N1 (“Mallard”) [64, 65], A/Udorn/307/1972 H3N2 (“Udorn”), A/England/195/2009 pH1N1 (“Eng195”), A/swine/England/87842/1990 H3N2 (“Swine H3N2”) and 2:6 reassortants of 2.3.4.4b H5N1 viruses containing PR8 segments 4 and 6 viruses were generated by reverse genetics as previously described[66]. Briefly, 2 × 10^6^ HEK293T cells were transfected in OptiMEM with pDUAL reverse genetics plasmids (250 ng of plasmid for each virus segment), using 4 µl of Lipofectamine 2000 according to the manufacturer’s instructions. At 24 hours post transfection, the media was replaced by serum-free DMEM supplemented with 0.14% (w/v) fraction V BSA and 1 μg/ml of l-(tosylamido-2-phenyl) ethyl chloromethyl ketone (TPCK)-treated trypsin. Virus-containing supernatant was collected after a further 2-days incubation (here described as a “P0 stock”). P0 stocks of PR8, Swine H3N2 and Udorn were propagated in MDCK cells. P0 stocks of Mallard, Eng195 and PR8 2:6 2.3.4.4b H5N1 reassortants were further propagated in 10-day embryonated hen’s eggs. Allantoic fluids were collected 48hpi. Cal04 P0 stocks were propagated in MDCK-SIAT cells. Reassortants between A/dairy cattle/Texas/24-008749-001-original/2024 H5N1 (“Cattle Texas”) and A/chicken/England/053052/2021 H5N1 (“AIV07”) were propagated in MDCK-GgANP32A.

**Table 1:**
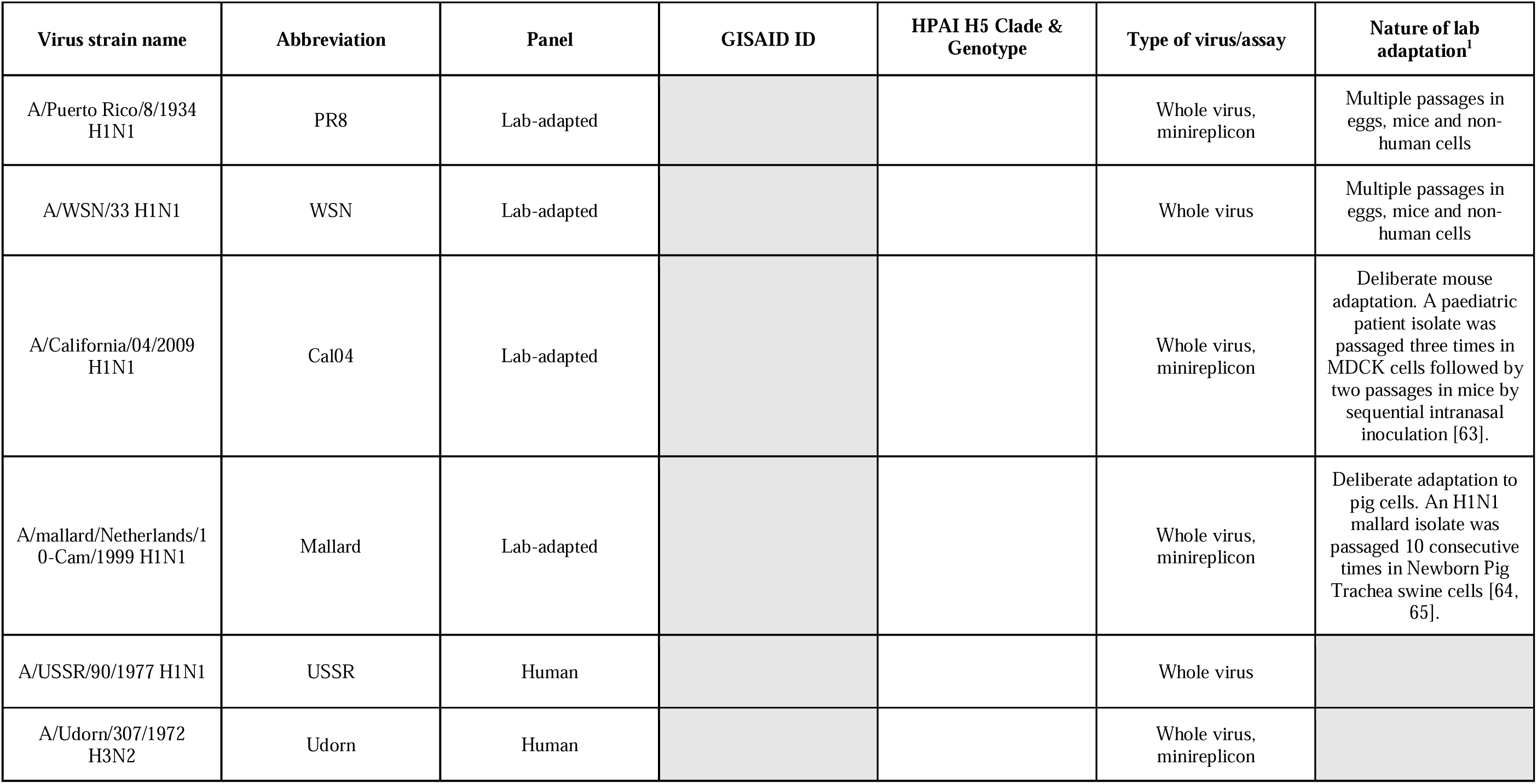
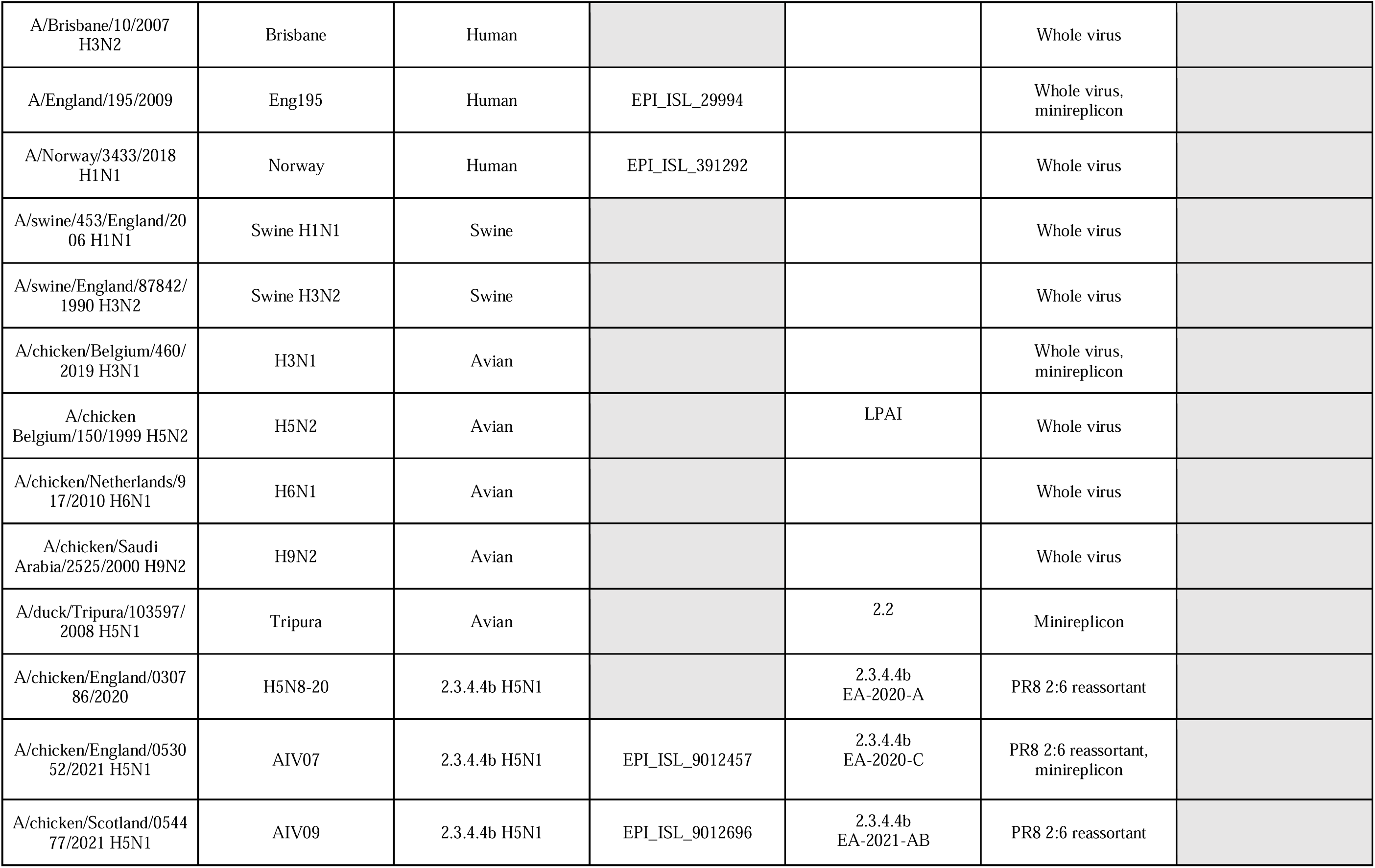
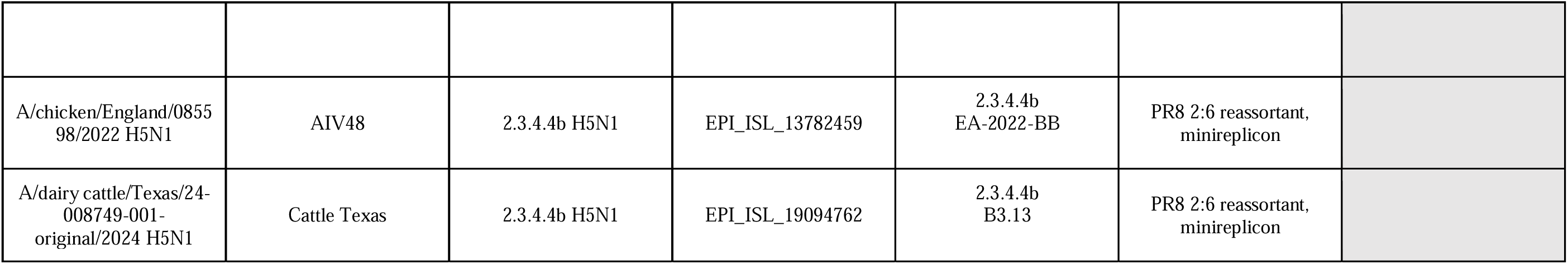
Virus strains, abbreviations and uses in this study. ^1^Laboratory adaptation was defined as viruses with a known history of passage in non-original host animals or cells and/or deliberate adaptation to a new host.

| Virus strain name | Abbreviation | Panel | GISAID ID | HPAI H5 Clade & Genotype | Type of virus/assay | Nature of lab adaptation<sup>1</sup> |
| --- | --- | --- | --- | --- | --- | --- |
| A/Puerto Rico/8/1934 H1N1 | PR8 | Lab-adapted |  |  | Whole virus, minireplicon | Multiple passages in eggs, mice and non-human cells |
| A/WSN/33 H1N1 | WSN | Lab-adapted |  |  | Whole virus | Multiple passages in eggs, mice and non-human cells |
| A/California/04/2009 H1N1 | Cal04 | Lab-adapted |  |  | Whole virus, minireplicon | Deliberate mouse adaptation. A paediatric patient isolate was passaged three times in MDCK cells followed by two passages in mice by sequential intranasal inoculation [63]. |
| A/mallard/Netherlands/10-Cam/1999 H1N1 | Mallard | Lab-adapted |  |  | Whole virus, minireplicon | Deliberate adaptation to pig cells. An H1N1 mallard isolate was passaged 10 consecutive times in Newborn Pig Trachea swine cells [64, 65]. |
| A/USSR/90/1977 H1N1 | USSR | Human |  |  | Whole virus |  |
| A/Udorn/307/1972 H3N2 | Udorn | Human |  |  | Whole virus, minireplicon |  |
| A/Brisbane/10/2007<br>H3N2 | Brisbane | Human |  |  | Whole virus |  |
| A/England/195/2009 | Eng195 | Human | EPI_ISL_29994 |  | Whole virus,<br>minireplicon |  |
| A/Norway/3433/2018<br>H1N1 | Norway | Human | EPI_ISL_391292 |  | Whole virus |  |
| A/swine/453/England/20<br>06 H1N1 | Swine H1N1 | Swine |  |  | Whole virus |  |
| A/swine/England/87842/<br>1990 H3N2 | Swine H3N2 | Swine |  |  | Whole virus |  |
| A/chicken/Belgium/460/<br>2019 H3N1 | H3N1 | Avian |  |  | Whole virus,<br>minireplicon |  |
| A/chicken<br>Belgium/150/1999 H5N2 | H5N2 | Avian |  | LPAI | Whole virus |  |
| A/chicken/Netherlands/9<br>17/2010 H6N1 | H6N1 | Avian |  |  | Whole virus |  |
| A/chicken/Saudi<br>Arabia/2525/2000 H9N2 | H9N2 | Avian |  |  | Whole virus |  |
| A/duck/Tripura/103597/<br>2008 H5N1 | Tripura | Avian |  | 2.2 | Minireplicon |  |
| A/chicken/England/0307<br>86/2020 | H5N8-20 | 2.3.4.4b H5N1 |  | 2.3.4.4b<br>EA-2020-A | PR8 2:6 reassortant |  |
| A/chicken/England/0530<br>52/2021 H5N1 | AIV07 | 2.3.4.4b H5N1 | EPI_ISL_9012457 | 2.3.4.4b<br>EA-2020-C | PR8 2:6 reassortant,<br>minireplicon |  |
| A/chicken/Scotland/0544<br>77/2021 H5N1 | AIV09 | 2.3.4.4b H5N1 | EPI_ISL_9012696 | 2.3.4.4b<br>EA-2021-AB | PR8 2:6 reassortant |  |
| A/chicken/England/0855<br>98/2022 H5N1 | AIV48 | 2.3.4.4b H5N1 | EPI_ISL_13782459 | 2.3.4.4b<br>EA-2022-BB | PR8 2:6 reassortant,<br>minireplicon |  |
| A/dairy cattle/Texas/24-008749-001-original/2024 H5N1 | Cattle Texas | 2.3.4.4b H5N1 | EPI_ISL_19094762 | 2.3.4.4b<br>B3.13 | PR8 2:6 reassortant,<br>minireplicon |  |

Isolates of A/USSR/90/1977 H1N1 (“USSR”) A/Brisbane/10/2007 H3N2 (“Brisbane”), A/Norway/3433/ 2018 H1N1 (“Norway”) and A/swine/453/England/2006 H1N1 (“Swine H1N1”) isolates were propagated in MDCK cells. Clarified supernatants were typically collected at 36–72 hpi. Isolates of A/chicken/Belgium/460/2019 H3N1 (“H3N1”), A/chicken Belgium/150/1999 H5N2 (“H5N2”), A/chicken/Netherlands/917/2010 H6N1 (“H6N1”) and A/chicken/Saudi Arabia/2525/2000 H9N2 (“H9N2”) isolates were propagated in hen’s eggs and allantoic fluids were collected and clarified at 48 hpi. All viruses were sequenced to confirm identity using Nanopore sequencing (described below).

### RNA extraction and Multi-segment RT-PCR

To confirm the genome sequences of the virus strains used, viral RNA was extracted from cell- and egg-grown viral stocks using the QIAamp Viral extraction mini kit (QIAGEN) as per manufacturer’s instructions. Viral genomes were amplified using the Superscript III One-Step RT-PCR kit (Invitrogen), 5 µl of extracted viral RNA, GGGGGGAGCAAAAGCAGG and GGGGGGAGCGAAAGCAGG as forward and GGGGTTATTAGTAGAAACAAGG as reverse primers[67]. Thermocycling conditions for RT-PCR reaction were: 55°C for 2 mins, 42°C for 60 mins, 94°C for 2 mins; 5 cycles of 94°C for 30 s, 44°C for 30 s and 68°C for 3 mins 30 s; 26 cycles of 94°C for 30 s, 57°C for 30 s and 68°C for 3 mins 30 s and 10 mins hold at 68°C. Resulting PCR was analysed on an 1% agarose gel.

### Nanopore sequencing and data analysis

RT-PCR products were pooled and prepared for sequencing using a ligation sequencing amplicon kit (SQK-NBD114.24, Oxford Nanopore Technologies, Oxford, UK) and sequenced on a FLO-MIN114 flow cell (R10.4.1., Oxford Nanopore Technologies) on an in-house GridION device. Final library concentration was approximately 10 fmol. Molarity was determined using Oxford Nanopore recommended tables, assuming an average multi-segment RT-PCR fragment size of 1.3 kb. Reads were called using the high accuracy basecall setting on the MinKNOW UI (Oxford Nanopore Technologies). The “passed” FASTQ output files were used in the sequencing analysis. Data was analysed using the EPI2ME (Oxford Nanopore Technologies) and Sequeduct (Edinburgh Genome Foundry) [68] pipelines. The processed FASTQ files were aligned against expected sequences. For the Sequeduct pipeline, reference sequences were converted to Genbank format. Consensus sequence output from the “analysis” workflow was used to align against expected sequences.

### Plaque assays

Confluent monolayers of MDCK cells (1 × 10^6^ cells seeded in 12-well plates the day prior to infection) were washed once with PBS, then infected with 250 µl of tenfold serial dilutions of virus. Following a 1 h incubation at 37 °C that allowed virus adsorption, cells were overlaid with DMEM supplemented with 0.14% fraction V BSA, 1 µg/ml TPCK-treated trypsin and 1.2% Avicel. Following a 48h undisturbed incubation at 37 °C, cells were fixed with 10% Neutral Buffered Formalin for at least 20 minutes and stained with a 0.1% toluidine blue solution for at least 1 h and plaques were then counted. Plaque assays for avian viruses or PR8 2:6 viruses with avian origin PB2 were performed as described but using MDCK-GgANP32A cells. Plaque assays for human viruses Eng195, Cal04, Brisbane and Norway H1N1 were performed using MDCK-SIAT cells and incubation temperatures of 35 °C for 72h.

### Virus replication kinetics

Monolayers of cells were washed once with PBS and infected with virus diluted in serum-free medium for 1 h at 37 °C (the multiplicity of infections used is specified in each figure legend). Infectious medium was replaced with serum-free medium supplemented with 0.14% fraction V BSA and 1 µg/ml TPCK-treated bovine pancreas trypsin.

Individual mammary explants were transferred to 96-well plates and infected with 5000 PFU of virus in 100 µl of serum free DMEM for 2h at 37°C. Explants were transferred to clean 24-well plates, washed 3x with warm PBS and overlayed with 1ml of DMEM supplemented with 0.14% fraction V BSA and 1µg/ml TPCK. Supernatant was collected at different timepoints post infection and infectious titres were determined by plaque assay.

### Immunoblotting

Cells were lysed in Laemmli buffer, heated at 80 °C for 5 min and subjected to polyacrylamide gel electrophoresis using sodium dodecyl sulfate-based buffer. Peptides were transferred to polyvinylidene difluoride membranes for 90 minutes at 32 V. Membranes were blocked for 1 h with 1× Tris-buffered saline with 0.1% Tween20 (TBST)/5% milk, washed three times with TBST and stained with primary antibodies diluted in TBST/5% BSA overnight at 4 °C. After three 5-minute TBST washes, secondary antibodies were diluted in blocking buffer and incubated for 1 h at room temperature protected from direct light. Following three more washes in TBST, membranes were imaged using the LI-COR Fc-Odyssey Imaging platform using the Image Studio Lite Software.

### Plasmid-based IFN- β promoter reporter assays

Under confluent monolayers of MAC-T or CLEC213 cells were co-transfected with 500ng or 100 ng of reporter plasmid encoding firefly luciferase under the control of human [69] or chicken IFN2 [70] promoter and 1000ng of segment 8 pDUAL plasmids using 2 μl of Lipofectamine 2000. 24 hours post transfection cells were stimulated with 5 μg of Poly I:C by transfection using 2 μl of Lipofectamine/well. Mock-stimulated cells were also included where Poly I:C was replaced by water. 24 hours post-stimulation, cells were lysed with 100 μl of reporter lysis buffer and luciferase activity was measured from 40 μl of lysate in opaque 96-well plates by using 25 μl of luciferase assay reagent in Cytation3 Plate Reader and Imager (integration time 1 second).

### Cellular gene expression shutoff transfection-based assays

Sub-confluent monolayers of MAC-T and CLEC213 cells were co-transfected in triplicate with a range of pDUAL segment 3 or 8 effector plasmids and 200ng of pRL using 2μL of Lipofectamine2000. Forty-eight hours post transfection, cells were lysed and *Renilla* luciferase expression was quantified using a *Renilla* Luciferase Assay kit. Briefly, cells were lysed with 100μL of 1× lysis buffer and frozen overnight at −20°C. Cells were scraped off the plate, cell debris was removed by centrifugation (8000rpm, 5 minutes, 4°C). Luminescence was measured from the mix of 20 μl of lysate and 50 μl of 1× *Renilla* luciferase substrate using a Cytation3 Plate Reader and Imager (integration time 1 second).

### Puromycin-labelling of nascent proteins

Subconfluent monolayers of MAC-T cells were infected at MOI 5. At 9, 12, 15 and 18 hpi, supernatant was replaced with fresh medium supplemented with 20 μg/ml of puromycin dihydrochloride for 1h. Cells were washed twice with PBS, lysed in Laemmli buffer and subjected to immunoblotting using a primary anti-puromycin antibody (Millipore; MABE343).

### RNP reconstitution (minireplicon) assays

Sub-confluent monolayers of MAC-T or CLEC213 cells (1 × 10^5^ cells seeded in 24-well plates the previous day) were co-transfected in triplicate with 50 ng of each pDUAL plasmid encoding PB2, PB1, PA and NP along with 20 ng of a bovine PolI-driven vRNA-like firefly luciferase reporter plasmid [17]. As negative controls, transfections lacking the PB2 plasmid (with empty pDUAL vector used to balance plasmid amount) were also performed. Two days post transfection, cells were lysed with 120 µl of reporter lysis buffer (Promega E4030). Lysates were cleared and luminescence was measured from 40 μl of lysate in opaque 96-well plates by using 25 μl of luciferase assay reagent in a Cytation3 Plate Reader and Imager (integration time 1 second).

### Tissue homogenisation

Mammary explants were added to 1ml of serum-free DMEM and a 5mm diameter stainless steel bead in 2ml tubes and homogenised using a Tissue Lyser II (Qiagen) system at 28kHz for 3 sets of 2 minutes. Homogenised tissue was clarified by centrifugation (3000 rpm, 5 min, 4°C). Homogenates were titrated for infectious virus by plaque assay.

### Glycomic analysis of MAC-T and bMEC cells

Approximately 10 million cultured MAC-T and bMEC cells were washed and scraped into cold PBS. Cells were then pelleted and washed several times with cold PBS prior to processing [71]. Briefly, cells were lysed by sonication, dialysed and lyophilised. Reduction and carboxymethylation reactions were carried out on the lyophilised material, which was subsequently dialysed and lyophilised. Samples were digested with trypsin (Sigma) and the resultant glycopeptides purified by Sep-Pak C18 chromatography. N-glycans were released by PNGase-F (Roche) and purified by Sep-Pak C18 chromatography. N-glycan samples were then split into three aliquots. Two of the three aliquots were treated with either sialidase-S or sialidase-A (Agilent Technologies) and then purified by C18 Sep-Pak chromatography, the third aliquot was untreated. Untreated and treated glycans were then permethylated and analysed using a 4800 MALDI-TOF mass spectrometer (Applied Biosystems).

MS and MS/MS data were visualised using Data Explorer (Applied Biosystems) and [M + Na]^+^ molecular ions were annotated manually with the assistance of GlycoWorkbench [72]. Glycan structures were predicted based on composition and knowledge of biosynthetic pathways, with MS/MS analysis used to confirm assignments. Figures were prepared using Adobe Illustrator.

Glycomic data was quantified using an R programme described in Wu et al., 2024 [73] which detects glycans according to their isotopic peak pattern and calculates the relative intensity across the isotope cluster. An R programme was then used to calculate the proportions of a-2,3 and a-2,6 sialic acid. First, the relative intensity of each glycan was normalised to the sum of the relative intensities of all the glycans in the sample. For sialylated glycans, normalised intensities were then weighted based on the number of sialic acid residues present on each glycan. The proportion of a-2,6 linked sialic acid was determined by summing the weighted normalised intensities of sialylated molecular ion peaks remaining after sialidase-S digestion. The proportion of a-2,3 linked sialic acid was calculated by subtracting the weighted normalised intensities of sialylated molecular ions in the sialidase-S sample from the weighted normalised intensities of sialylated molecular ions from the untreated sample (raw data in Supplementary Table 2) [74].

### Cell staining and flow cytometry

Infected bMECs were trypsinised, resuspended and fixed in PBS/4% formaldehyde. Following a PBS wash by centrifugation, cells were blocked with PBS/2%BSA for 1h. Cells were stained with rabbit anti-H3 (c33) or mouse anti-H1 (2-12C, Absolute antibody) primary antibodies diluted 1:5000 in blocking buffer for 1h at room temperature and with constant agitation. Cells were then washed twice 2x with blocking buffer and resuspended in a secondary antibody solution containing donkey anti-rabbit647 (ab150075, abcam) and donkey anti-mouse 488 (A11011, Invitrogen) for another 30 min. Cells were centrifuged, washed twice with blocking buffer and analysed using a BD LSR Fortessa X20 cytometer. The gating strategy used to define the infected cell populations is shown in Figure S13.

## Acknowledgments

The authors would like to thank Prof Massimo Palmarini (MRC-University of Glasgow Centre for Virus Research) and Dr John McCauley (recently retired from Crick Institute) for providing MDCK-GgANP32A and MDCK-SIAT cells, respectively. BAT-II cells were kindly provided by Dr Diane Lee (University of Surrey). Chicken lung cells (CLEC213) were kindly provided by Dr Sascha Trapp of the INRA Centre Val de Loire. We thank Dr Laurence Tiley (The University of Cambridge), Dr Ron Fouchier (Erasmus MC) and Dr Daniel Perez (The University of Georgia) for providing the reverse genetics systems for A/mallard/Netherlands/10-Cam/1999 (H1N1), A/Puerto Rico/8/1934 (H1N1) and A/California/04-061-MA/2009 (H1N1) viruses, respectively, as well as colleagues at the Animal and Plant Health Agency for early provision of viral genetic data to the FluMap consortium. Human IFN-β::firefly luciferase and chicken IFN2::firefly luciferase reporter plasmids were kindly gifted by Prof Richard Randall (University of St Andrews) and Dr Holly Shelton (previously at Pirbright Institute), respectively. We gratefully acknowledge the expert assistance of staff at the University of Edinburgh Large Animal Research Imaging Facility and Langhill Farm. For the purpose of open access, the author has applied a Creative Commons Attribution (CC BY) licence to any Author Accepted Manuscript version arising from this submission.

## Funding

UK Biotechnology and Biological Sciences Research Council grant BB/CCG2270/1 (The Roslin Immunological Toolbox)

UK Biotechnology and Biological Sciences Research Council Institute Strategic Programme funding grants BBS/E/RL/230002A and BBS/E/RL/230002C (RMP, JRF, JH, JS, TB, KS, FXD, EG and PD)

UK Biotechnology and Biological Sciences Research Council Institute Strategic Programme funding grant BBS/E/PI/230002B and National Bioscience Research Infrastructure funding BBS/E/PI/23NB0004 and BBS/E/PI/23NB0003 (TPP, IB and MI)

UK Biotechnology and Biological Sciences Research Council grant BB/V004697/1 (PRM)

UK Biotechnology and Biological Sciences Research Council grant BB/V011286/1 (PD)

UK Natural Environment Research Council (NERC) Capability to Deliver funding NE/Y001591/1 (EG and PD)

UK Medical Research Council FluTrailMAP-One Health consortium grant MR/Y03368X/1 (RMP, TPP, IB, WB, DHG, MI, PRM, SMH and PD

UK Medical Research Council grant MR/Y015045/1 (PD)

UK Biotechnology and Biological Sciences Research Council and UK Department for Environment, Food & Rural Affairs FluMap consortium grants BB/X006123/1 and BB/X006166/1 (PD, MI)

UK Biotechnology and Biological Sciences Research Council and UK Department for Environment, Food & Rural Affairs FluTrailmap consortium grants BB/Y007352/1 and BB/Y007298/1 (RMP, EG, PD, TPP, IB, MI)

UK Biotechnology and Biological Sciences Research Council and Department of Biotechnology, Government of India grants BB/L004666/1 and BT/IN/Indo-UK/FADH/48/AM/2013 (AAR, AM and PD).

UK Biotechnology and Biological Sciences Research Council and Department of Biotechnology and ERA-net ICRAD project FluNuance grant number BB/V0119899/1 (SdW, KS and PD)

Horizon Europe project 101287378 (EUPAHW2.0) (RMP, JRF, EG and PD). Views and opinions expressed are however those of the author(s) only and do not necessarily reflect those of the European Union or the European Research Executive Agency. Neither the European Union nor the granting authority can be held responsible for them.

Wellcome Trust Sir Henry Dale fellowship award 211222/Z/18/Z (EG).

UK Academy of Medical Sciences Springboard Grant 1049 (DHG)

University of Edinburgh Chancellor’s Fellowship (RMP)

Roslin Foundation PhD studentship (MB)

UK Biotechnology and Biological Sciences Research Council EastBio Doctoral Training Partnership

PhD studentship BB/T00875X/1 (AM).

## Credit contributions

Rute Maria Pinto – RMP – Conceptualisation, Methodology, Validation, Formal analysis, Investigation, Resources, Data curation, Visualisation, Writing – original draft, Writing – reviewing and editing, Supervision, Project administration, Funding acquisition

Colin P Sharp – CS -Conceptualisation, Methodology, Validation, Investigation, Resources, Data curation, Writing – reviewing and editing, Supervision, Project administration

Maia Beeson – MB – Methodology, Validation, Investigation, Data curation, Writing – reviewing and editing

Nunticha Pankaew – NP – Methodology, Validation, Investigation, Data curation, Writing – reviewing and editing

Jack A. Hassard - JAH – Methodology, Validation, Formal analysis, Investigation, Data curation, Visualisation, Writing – original draft, Writing – reviewing and editing

Alexander Moxom – AM – Methodology, Validation, Investigation, Writing – original draft, Writing – reviewing and editing

Callum Magill – CM – Methodology, Validation, Investigation, Writing – original draft

Rebecca A Ross – RAR- Methodology, Validation, Investigation, Writing - reviewing and editing

Laura Tuck – LT – Methodology, Validation, Writing – reviewing and editing

Stephen Meek – SM – Methodology, Writing – reviewing and editing

Hui Min Lee – HML – Methodology, Resources

Kirsty Jensen – JK – Methodology, Writing – reviewing and editing

Inga Dry – ID – Methodology, Validation, Writing – reviewing and editing

Pedro Melo – PM – Methodology, Resources, Writing – reviewing and editing

Wenfang Spring Tan – WST – Methodology, Investigation

Jiayun Yang – JY.- Methodology, Resources, Writing – reviewing and editing

Ashwin Ashok Raut – AAR - Resources

Anamika Mishra – AM - Resources

Sjaak de Wit – SdW – Resources, Writing – reviewing and editing

J. Ross Fitzgerald – JRF – Methodology, Resources, Writing – reviewing and editing, Funding acquisition

Jayne C. Hope – JCH – Methodology, Resources, Writing – reviewing and editing, Funding acquisition

Joanne Stevens – JS – Methodology, Resources, Funding acquisition

Tom Burdon – TB – Methodology, Resources, Funding acquisition

Kate Sutton – KS – Methodology, Resources, Writing – reviewing and editing, Funding acquisition

Cristina L. Esteves – CLE – Methodology, Resources

F. Xavier Donadeu – FXD – Methodology, Resources, Writing – reviewing and editing, Funding acquisition

Thomas P. Peacock – TPP – Methodology, Writing – reviewing and editing, Funding acquisition.

Ian Brown - IB - Project administration, Funding acquisition

Wendy Barclay – WB - Project administration, Funding acquisition

Daniel H. Goldhill – DHG – Methodology, Resources, Writing – reviewing and editing, Funding acquisition

Munir Iqbal – MI – Methodology, Resources, Writing – reviewing and editing, Funding acquisition

Pablo R. Murcia – PMR – Methodology, Resources, Writing – reviewing and editing, Funding acquisition

Stuart M. Haslam – SMH – Methodology, Resources, Supervision, Project administration, Funding acquisition

Eleanor Gaunt – EG – Conceptualisation, Methodology, Resources, Data curation, Visualisation, Writing – original draft, Writing – reviewing and editing, Supervision, Project administration, Funding acquisition

Paul Digard – PD – Conceptualisation, Methodology, Resources, Data curation, Writing – original draft, Writing – reviewing and editing, Supervision, Project administration, Funding acquisition

## Data and materials availability

All data and materials used in the analysis will be available upon reasonable request. Some reagents will require a material transfer agreement (MTA).

---

**Figure S1:**
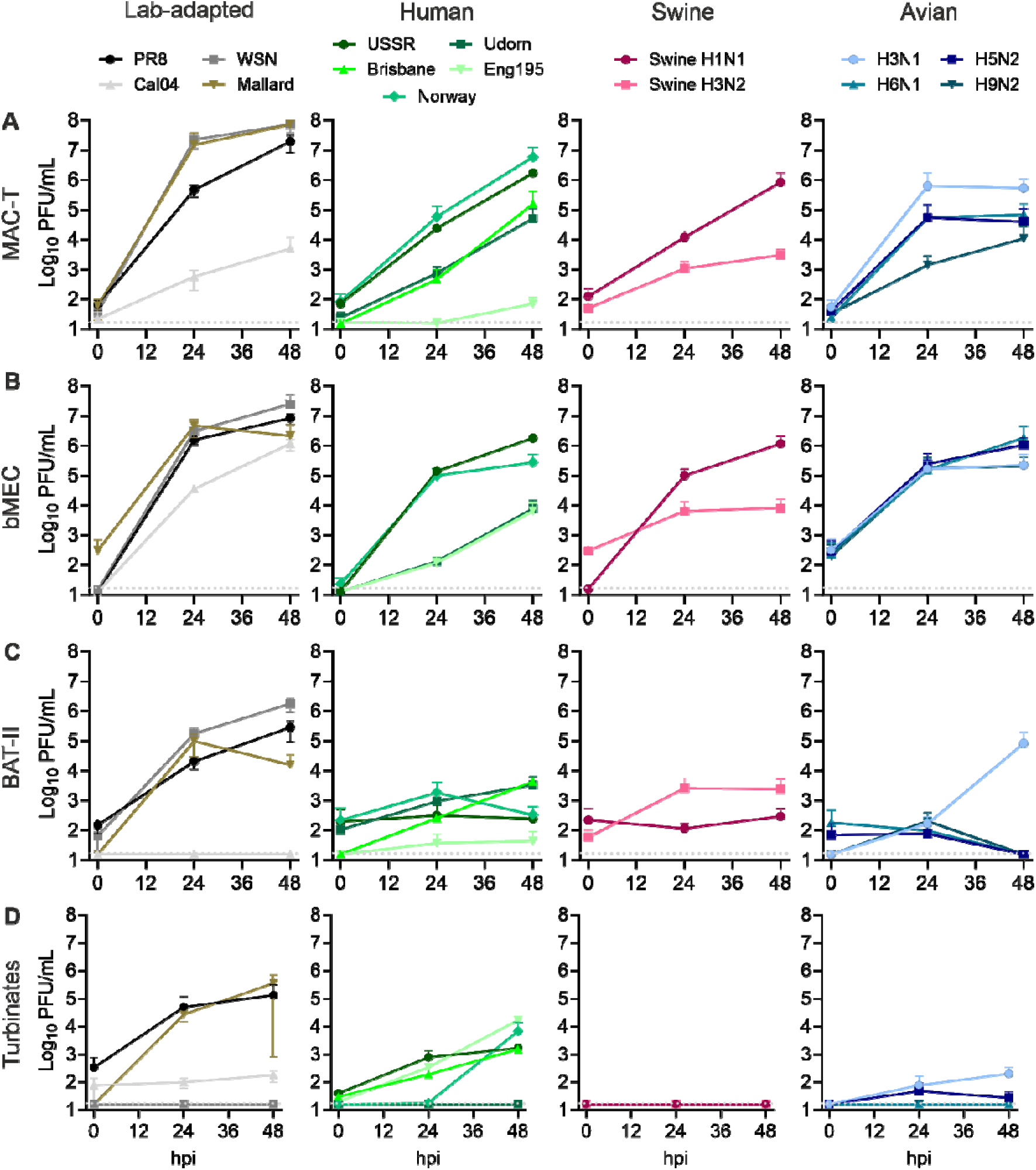
IAV replication kinetics in cow mammary and respiratory cells. MAC-T (A), bMEC (B), BAT-II (C) and nasal turbinates (D) were infected with the indicated viruses at MOI 0.1 (for bMEC) or 0.01 (for the remaining cells) and supernatants were collected at 0, 24 and 48 hpi. Viral titres were determined by plaque assay. Data represent mean ± SEM from three independent experiments each performed in duplicate.

**Figure S2:**
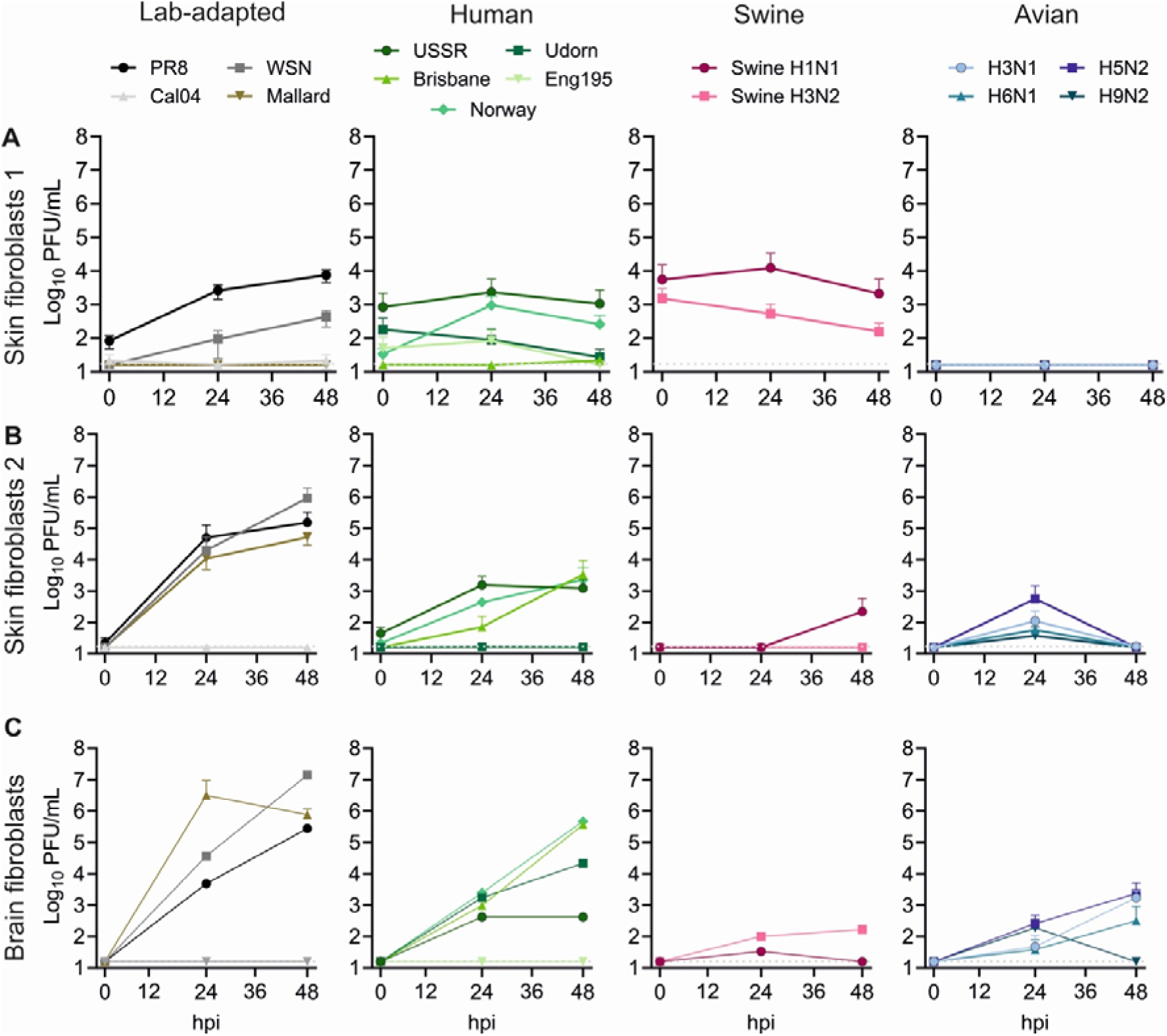
IAV replication kinetics in cow fibroblasts. Skin 1 (A), skin 2 (B) and brain (C) primary fibroblasts were infected with the indicated viruses at MOI 0.01 and supernatants were collected at 0, 24 and 48 hpi. Viral titres were determined by plaque assay. Data represent mean ± SEM from three independent experiments each performed in duplicate.

**Figure S3:**
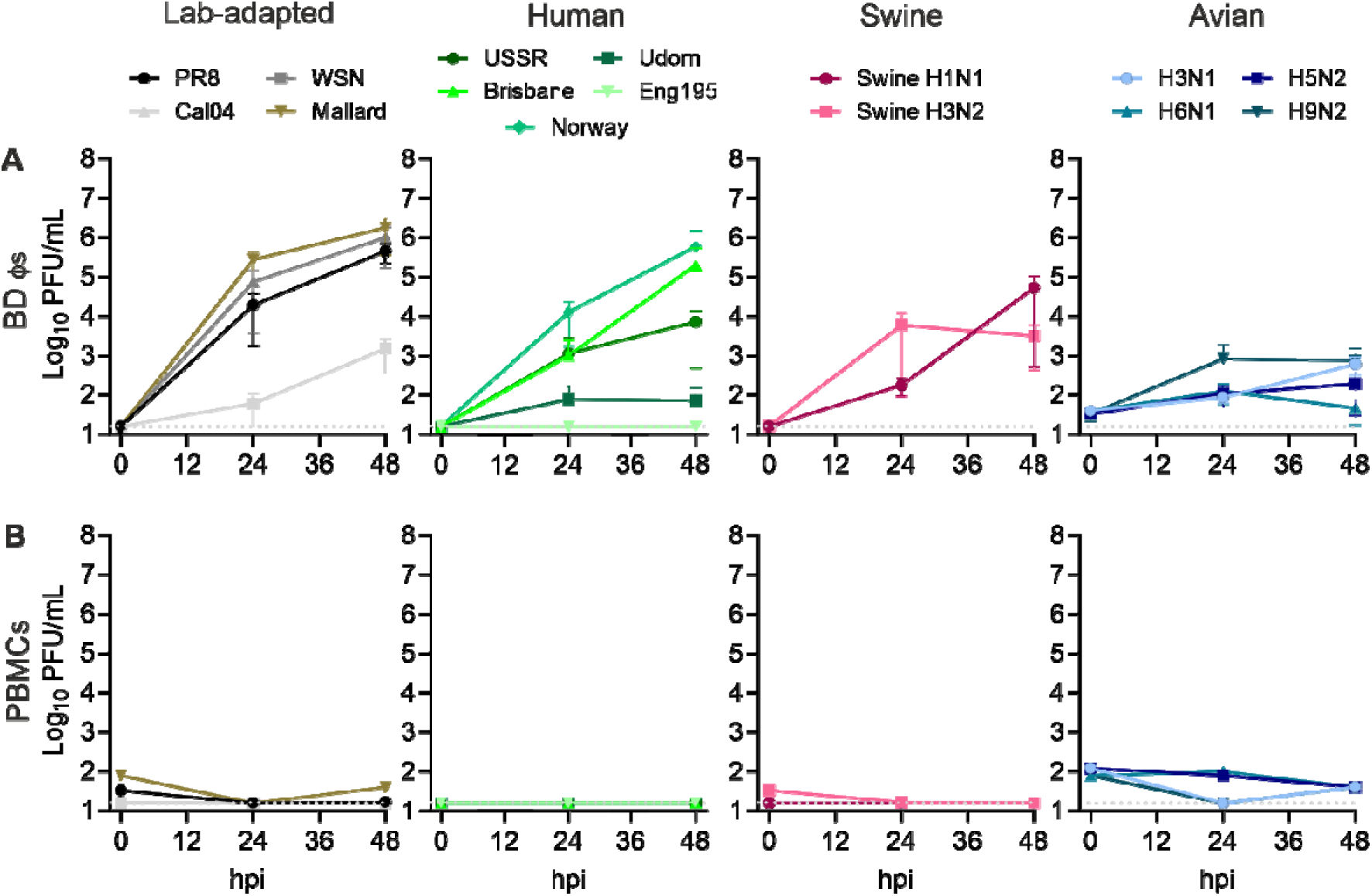
IAV replication kinetics in immune cells. Blood-derived macrophages (BD□) (A) or peripheral blood mononuclear cells (PBMC) (B) were infected with the indicated viruses at MOI 0.01 and supernatants were collected at 0, 24 and 48 hpi. Viral titres were determined by plaque assay. Data represent mean ± SEM from three independent (BD s) or a single (PBMCs) experiment each performed in duplicate.

**Figure S4:**
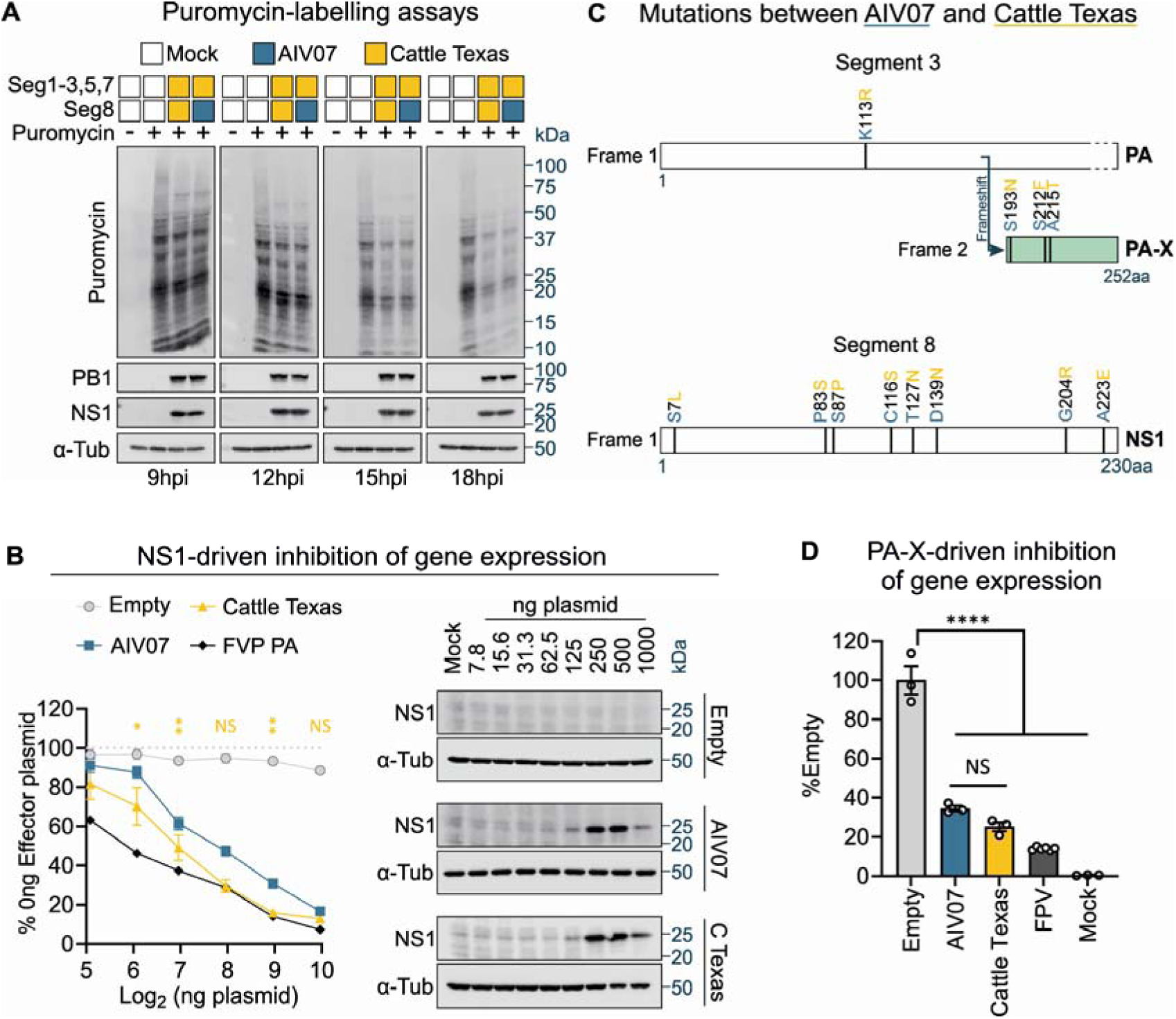
Cattle Texas host shutoff activity is similar to that of AIV07. (A) MAC-T cells were infected at MOI 5 and pulsed with puromycin at the indicated times post infection for 1h. Cells were lysed and new protein synthesis was measured by western blotting for puromycin. NS1 and PB1 were detected to validate infection and α-Tubulin as a loading control. The positions of molecular weight markers are indicated. (B) MAC-T cells were transfected with a reporter plasmid constitutively expressing *Renilla* luciferase under the control of a CMV promoter alongside a dose range of pDUAL plasmids expressing AIV07 or Cattle Texas NS1s. A dilution series of pDUAL plasmid expressing segment 3 of A/Chicken/Rostock/1934 H7N1 (FPV) was used as a positive control. Data represent the mean ± SEM of three independent experiments each performed in triplicate. Statistical annotations are the result of one-way ANOVA analysis performed for each plasmid dosage. Shown multiple comparisons were made between AIV07 and Cattle Texas using a Dunnett’s test at the given segment 8 dosages. Immunoblotting was performed to detect increasing levels of PB1 and NS1 expression, using α-Tubulin as loading control. (C) Schematic representation of PA-X and NS1 mutations between AIV07 and Cattle Texas. Open reading frames from frames 1 and 2 are represented by white and mint boxes, respectively. Amino acid differences between AIV07 (blue) and Cattle Texas in PA-X and NS1 are highlighted. (D) MAC-T cells were transfected with a reporter plasmid constitutively expressing *Renilla* luciferase under the control of a CMV promoter alongside segment 3 pDUAL plasmids (expressing PA-X). Data represent mean ± SEM from three independent experiments each performed in triplicate. Statistical annotations are the result of one-way ANOVA analysis and multiple comparisons performed using a Dunnett’s test. For (B) and (D) Luminescence readings were measured 48h post transfection. For all shown statistical annotations: NS indicates non significant, * a p-value <0.05, ** a p-value <0.01, *** a p-value <0.001 and **** a p-value <0.0001.

**Figure S5:**
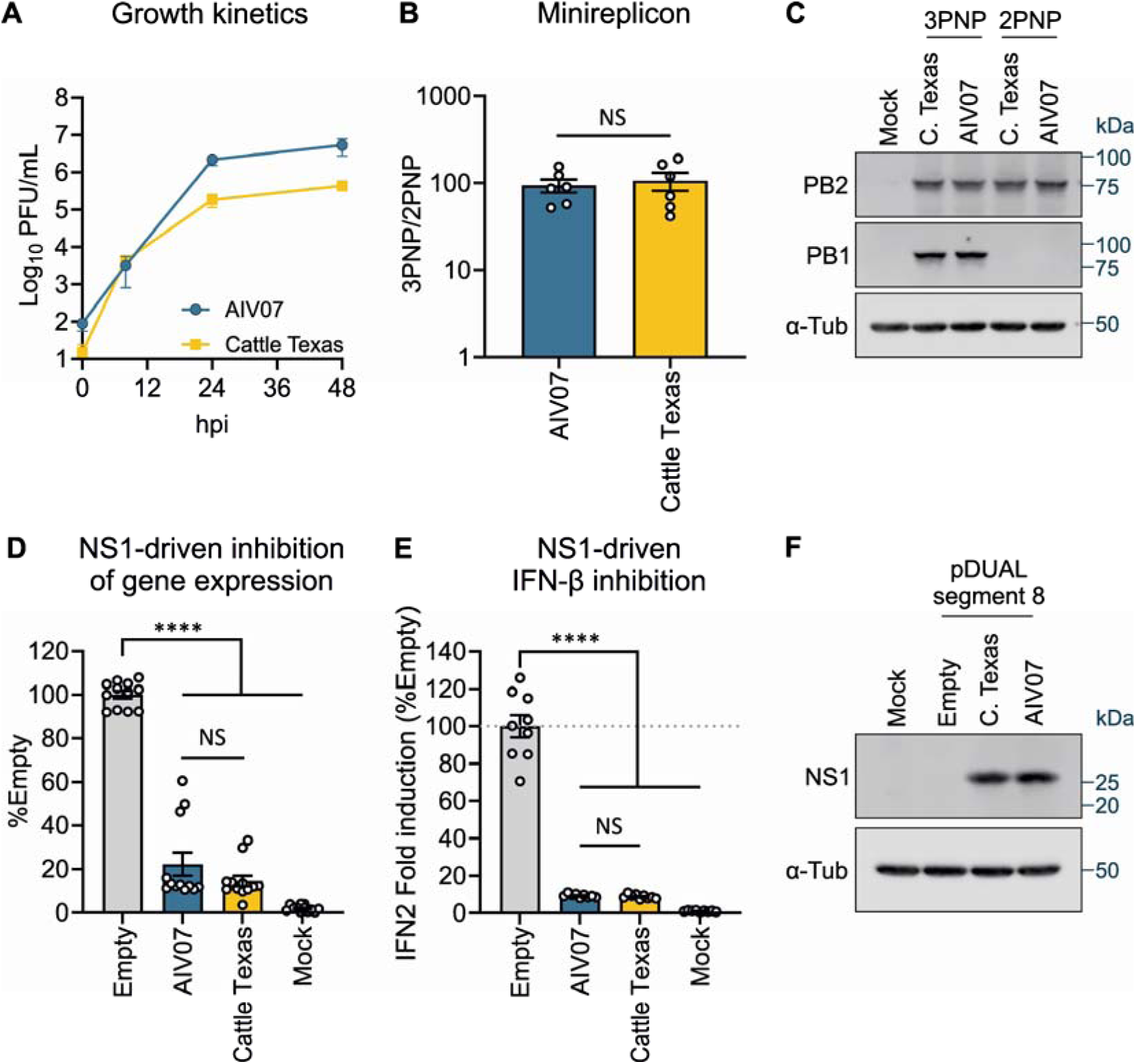
Cattle Texas is not fitter in chicken cells. (A) CLEC213 chicken lung epithelial cells were infected with the indicated viruses at MOI 0.01. Supernatants were collected at the indicated times post infection and virus titres were determined by plaque assay. Data represent mean ± SEM from three independent experiments each performed in duplicate. (B) CLEC213 cells were transfected with pDUAL plasmids encoding segments 1-3 and 5 of the indicated combinations of AIV07 and Cattle Texas RNP complexes alongside a firefly luciferase-expressing vRNA-like reporter plasmid. Luminescence levels were measured at 48h post transfection. Data represent mean ± SEM from three independent experiments each performed in duplicate. Statistical annotations are the result of an unpaired t-test. (C) Expression of polymerase components from (B) was examined by western blot. α-Tubulin was used as loading control. The positions of molecular weight markers are indicated. (D) CLEC213 cells were transfected with a reporter plasmid constitutively expressing *Renilla* luciferase under the control of a CMV promoter alongside AIV07 or Cattle Texas segment 8 or empty vector plasmids and luminescence readings measured 48h later. Data represent mean ± SEM from four independent experiments each performed in triplicate. Statistical annotations are the result of one-way ANOVA analysis. (E) CLEC213 cells were transfected with a chicken IFN2 promoter-driven firefly luciferase reporter alongside AIV07 or Cattle Texas segment 8 plasmids, and poly I:C treated 24h later. Luminescence readings were acquired after a further 24h. Data represent mean ± SEM from three independent experiments each performed in duplicate. Statistical annotations are the result of one-way ANOVA analysis. (F) NS1 expression from (A) and (B) was tested by western blot. α-Tubulin was used as loading control. For D and F, multiple comparisons were performed using a Dunnett’s test. For all shown statistical annotations: NS – non significant and ****p-value <0.0001.

**Figure S6:**
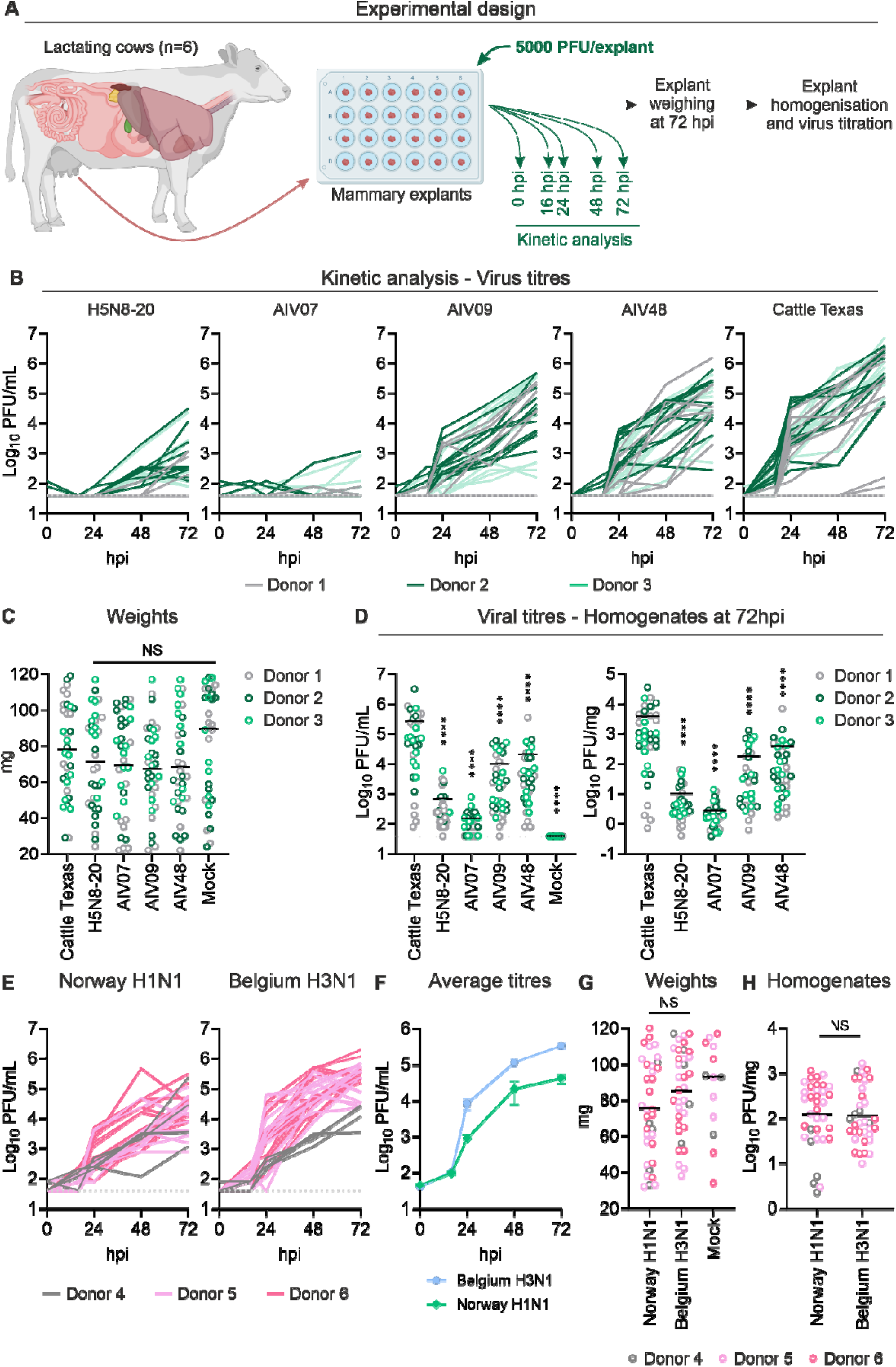
Infections in bovine mammary explants. (A) Schematic representation of the experimental design. (B) Multiple explants were extracted from the mammary glands from 3 donors and infected with 5000 PFU/explant of the indicated PR8 2:6 reassortant viruses. Supernatants were collected at the indicated times post infection and virus titres determined by plaque assay. Data are from individual explants coloured by donor: donor 1 (n=12), donor 2 (n=12), donor 3 (n=10). (C) Weights of explants from B at 72 hpi. (D) Explants were homogenised, viral titres measured by plaque assay (left panel) and normalised to individual explant weights (right panel). For C-D, black lines indicate means, circles indicate data for individual explants colour-coded by donor. Statistical annotations are the result of a one-way ANOVA test. Statistical comparisons are given against Cattle Texas using a Dunnet’s test. (E) Explants were extracted from the mammary glands from 3 donors and infected with 5000 PFU/explant of Norway of Belgium H3N1 viruses. Supernatants were collected at the indicated times post infection and virus titres determined by plaque assay. Data are from individual explants coloured by donor: donor 4 (n=5), donor 5 (n=15), donor 6 (n=15). (F). Averages from E. (G) Weights of explants from E at 72 hpi. Statistical annotations are the result of a one-way ANOVA test. Comparisons were performed using a Dunnet’s test. (H) Explants were homogenised, viral titres measured by plaque assay and normalised to individual explant weights. Statistical annotations are the result of a t-test. For G and H, black lines indicate means, circles indicate data for individual explants colour-coded by donor. For all shown statistical annotations: *p-value <0.05, **p-value <0.01, ***p-value <0.001, ****p-value <0.0001, NS – non-significant.

**Figure S7:**
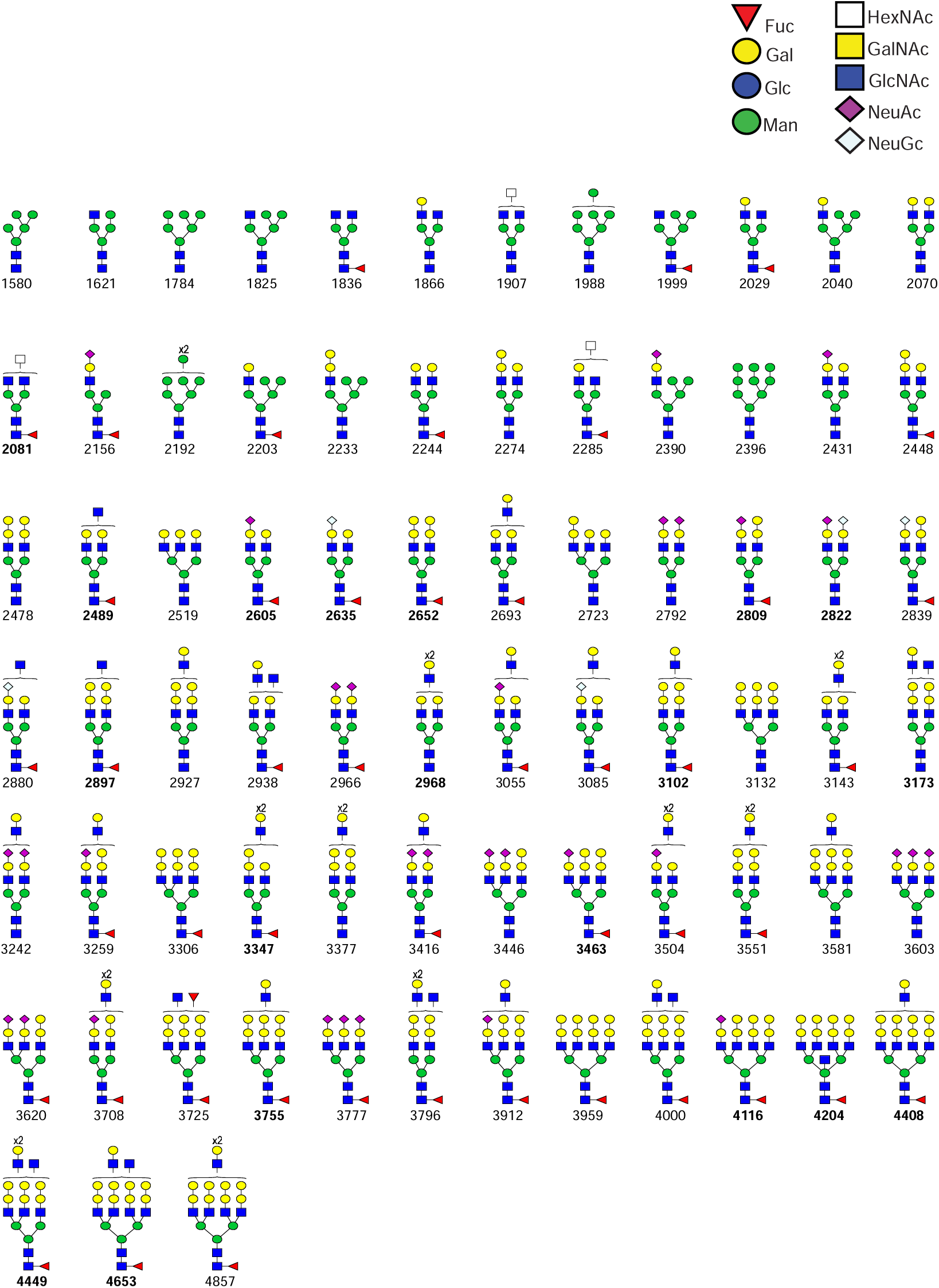
Structures of N-glycans from MAC-T cells. Proposed structures of the N-glycans from MAC-T cells. Structures were based on composition, biosynthetic knowledge and MS/MS analysis. Structures in bold have been confirmed directly by MS/MS. Structures were drawn according to the Symbol Nomenclature for Glycans (SNFG) guidelines.

**Figure S8:**
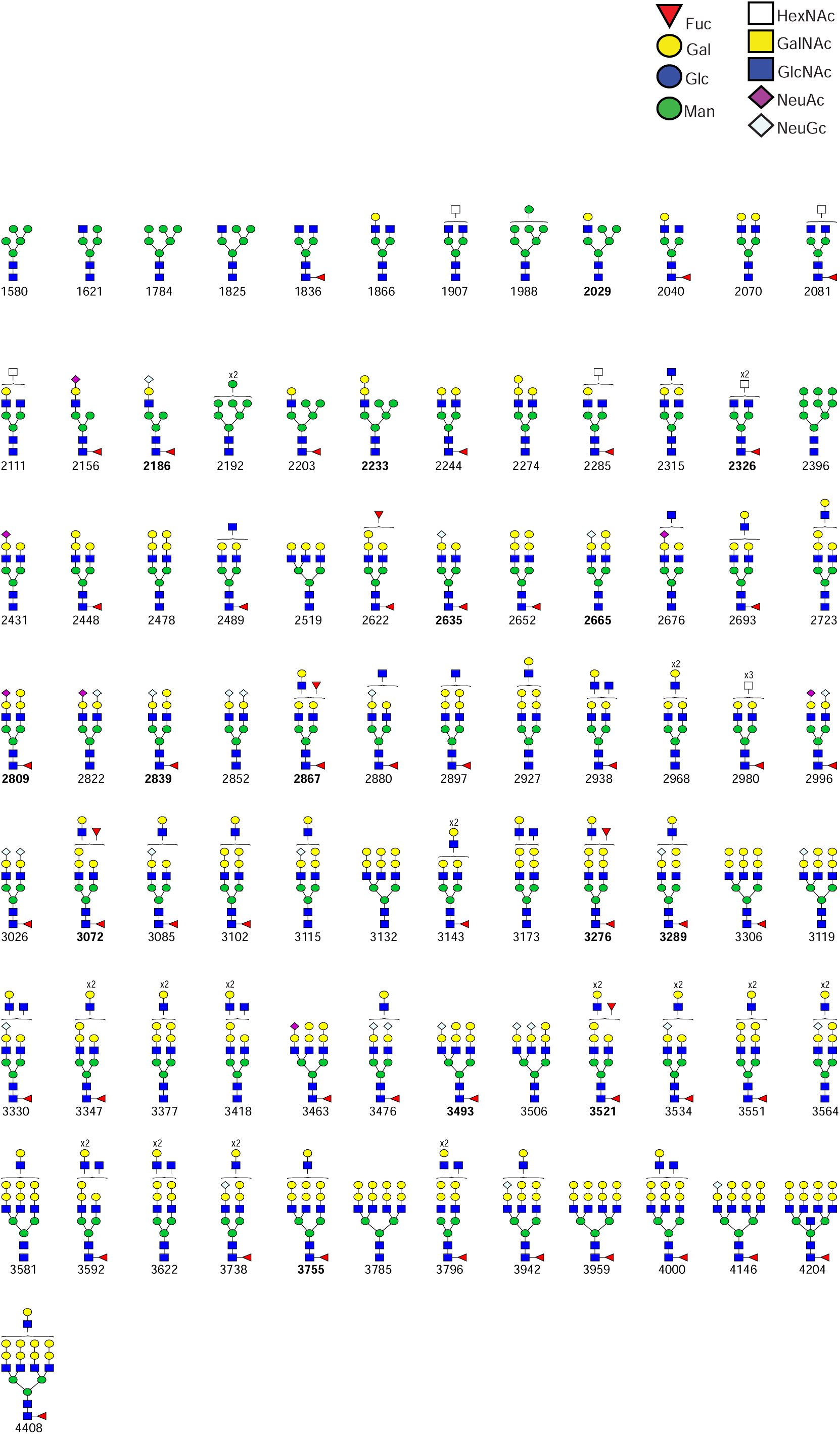
Structures of N-glycans from bMEC cells. Proposed structures of the N-glycans from bMEC cells. Structures were based on composition, biosynthetic knowledge and MS/MS analysis. Structures in bold have been confirmed directly by MS/MS. Structures were drawn according to the Symbol Nomenclature for Glycans (SNFG) guidelines.

**Figure S9:**
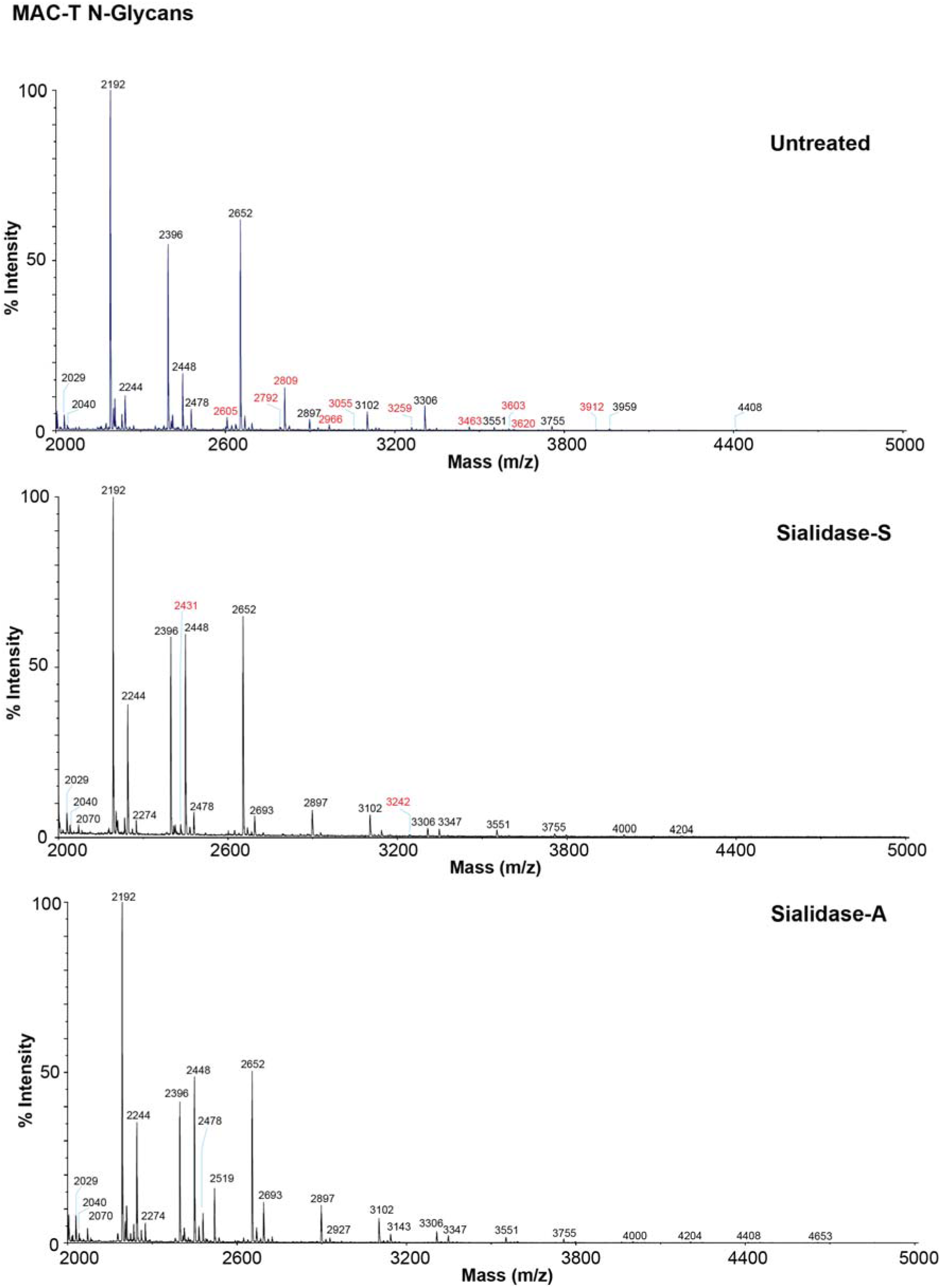
MALDI-TOF spectra of permethylated N-glycans from MAC-T cells following sialidase digestion. MALDI-TOF spectra of permethylated N-glycans from MAC-T cells treated with sialidase-S or sialidase-A. Annotations show [M + Na]^+^ molecular ions and include major structures which were unsialylated (black) and sialylated (red).

**Figure S10:**
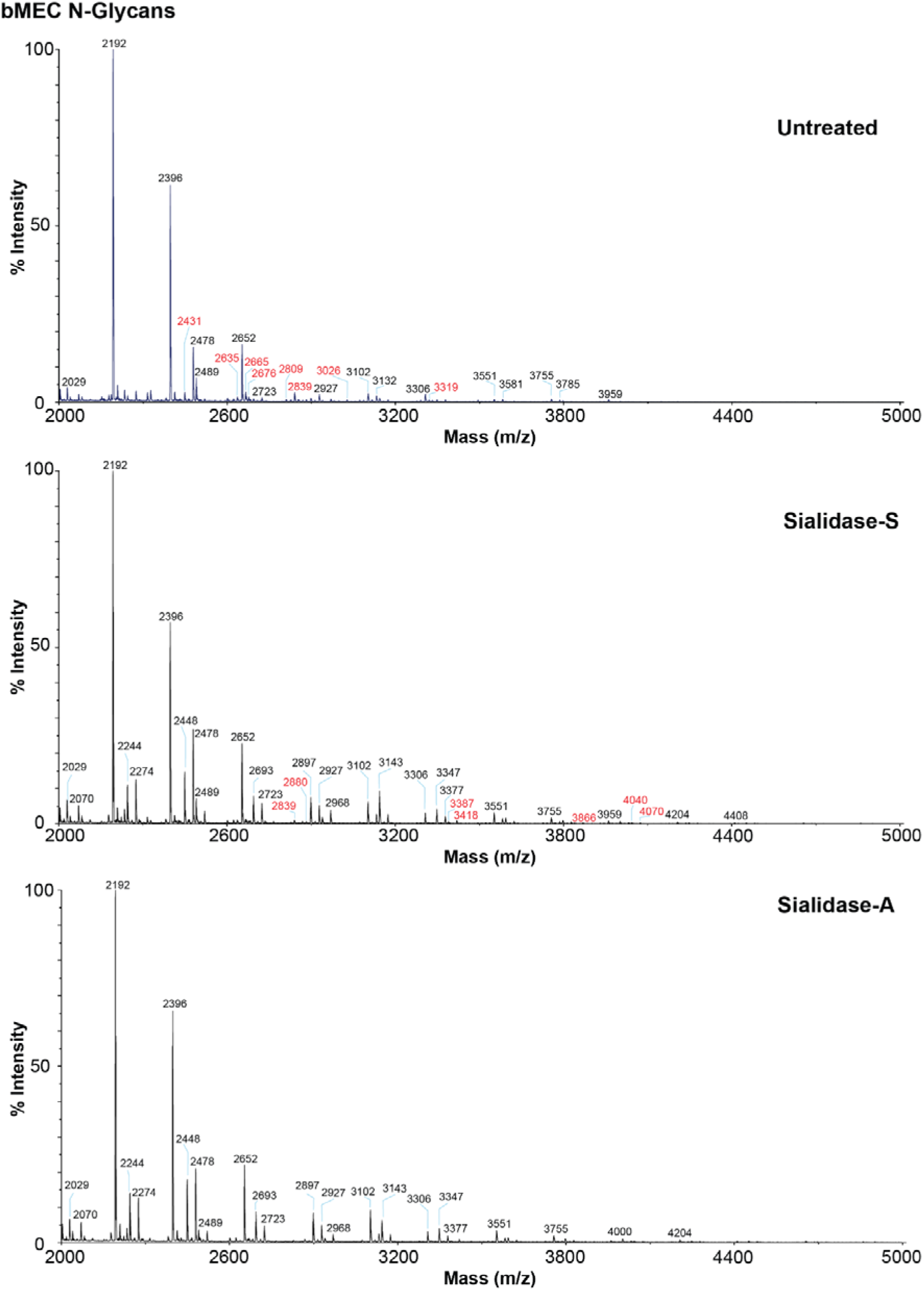
MALDI-TOF spectra of permethylated N-glycans from bMEC cells following sialidase digestion. MALDI-TOF spectra of permethylated N-glycans from MAC-T cells treated with sialidase-S or sialidase-A. Annotations show [M + Na]^+^ molecular ions and include major structures which were unsialylated (black) and sialylated (red).

**Figure S11:**
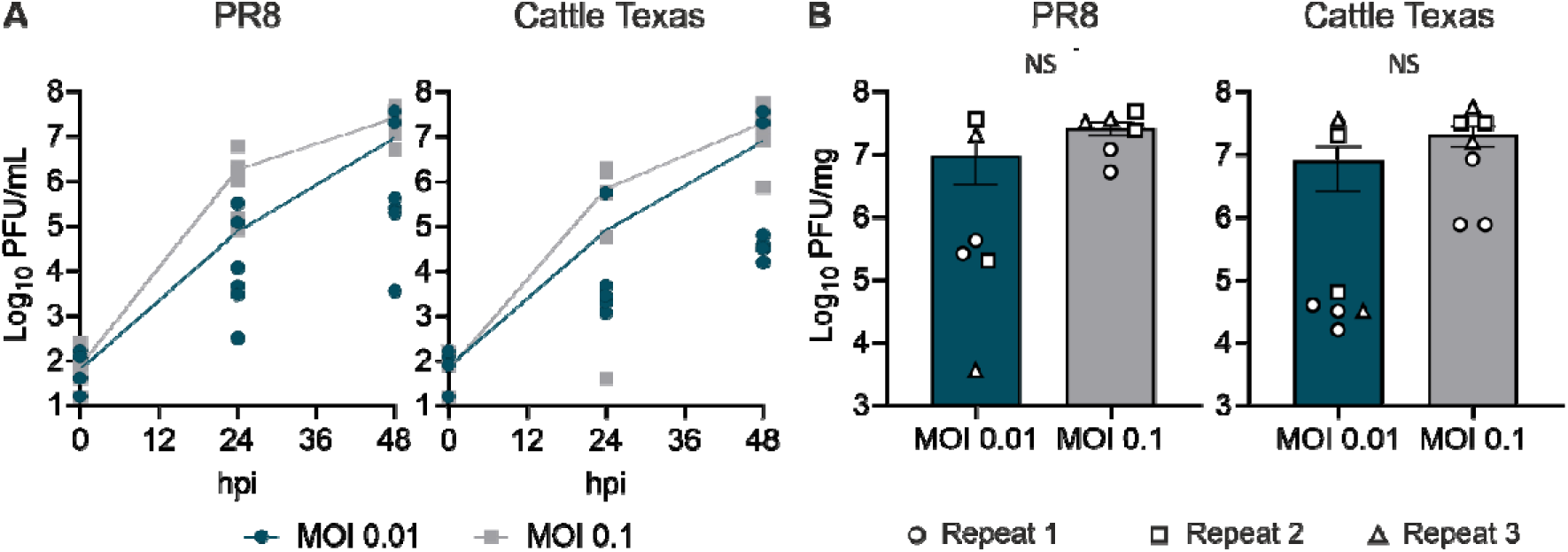
Optimisation of bMEC multicycle infection. A) bMECs were infected with PR8 or PR8 2:6 reassortant of Cattle Texas (here determined Cattle Texas) at MOI 0.01 or 0.1. Supernatants were collected at 0, 24 and 48 hpi and viral titres were determined by plaque assay. Data represent mean ± SEM from three independent experiments each performed in duplicate or triplicate. B) End point titres from A are shown. Individual technical replicates from the 3 different repeats are shown. Statistical annotations are the result of an unpaired t test. NS – non-significant.

**Figure S12:**
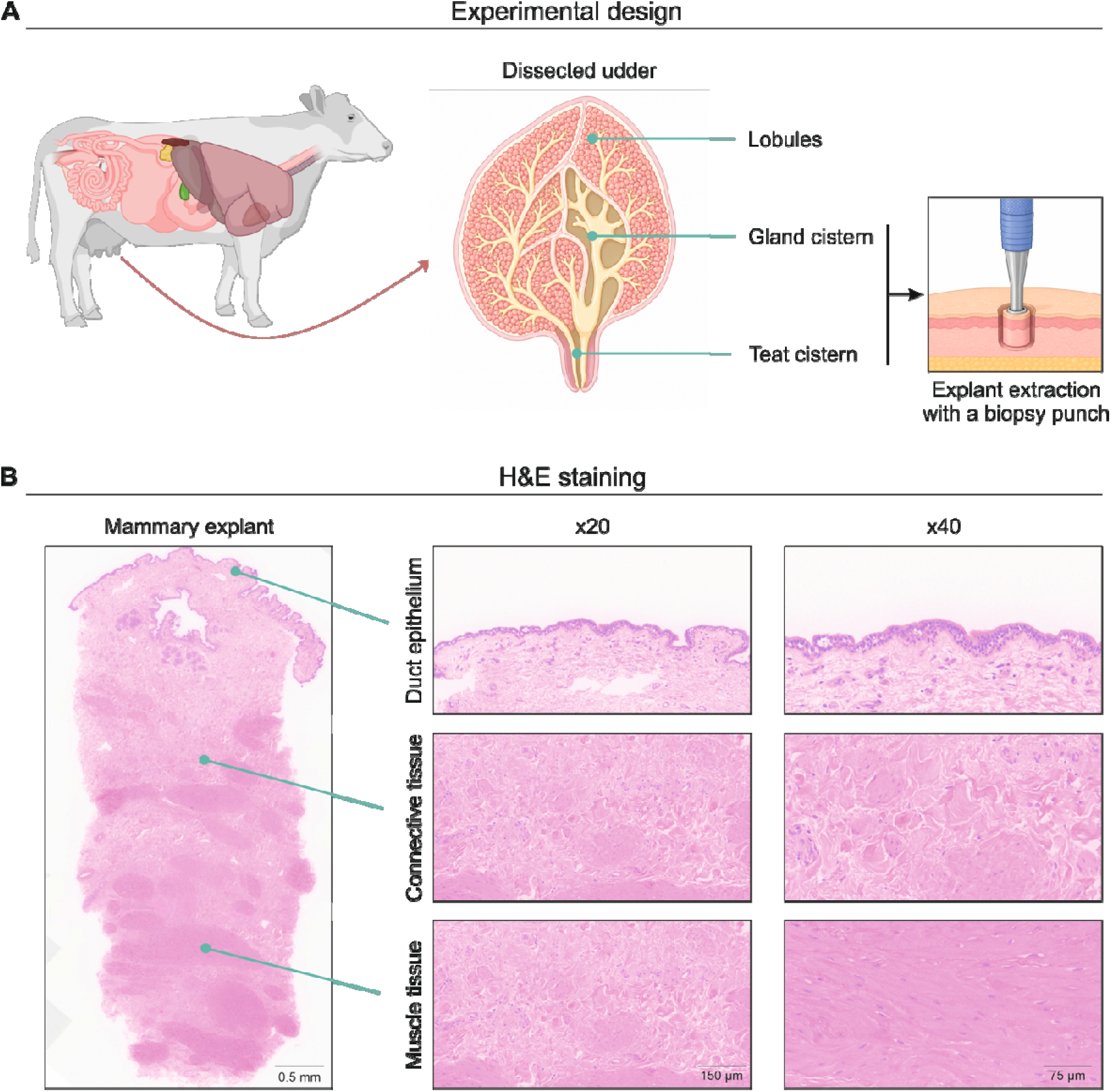
Histopathology of bovine mammary explants. (A) Schematic representation of extraction of mammary explants. (B) Representative figures of H&E staining and histopathology performed in bovine mammary explants extracted from the gland or teat cisterns.

**Figure S13:**
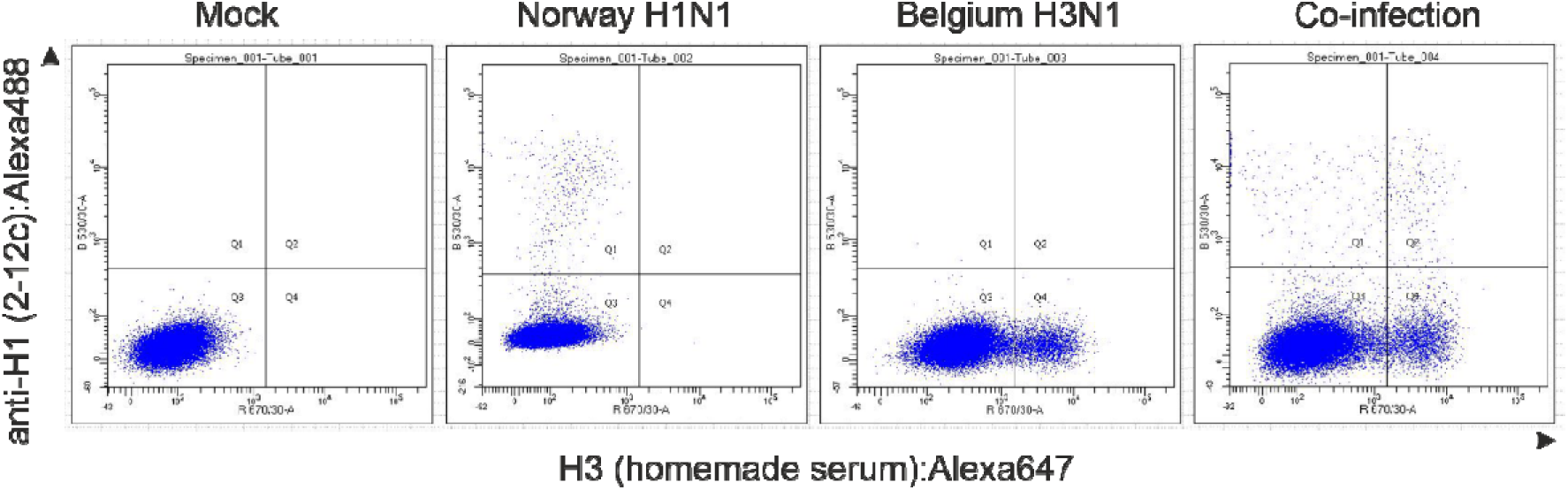
Flow cytometry gating strategy for examining co-infection of bovine mammary gland cells. bMECs were infected with human virus Norway H1N1 or/and chicken virus Belgium H3N1 for 16h at MOI 1. Cells were stained with anti-H1 and H3 antibodies and analysed by flow cytometry. Figure shows a gating example.

**Table S1.** Amino acid differences between the internal gene products of avian precursor AIV07 isolate and bovine isolate B3.13. Complete genome sequences were downloaded from GISAID (EPI_ISL_9012457 and EPI_ISL_19014384 respectively), aligned in SSEv1.4 sequence editing software and annotated manually.

| Protein | Amino acid changes |
| --- | --- |
| PB2 | T58A G79S V109I V139I F154L E362G D441N V478I V495I M631L V649I T676A |
| PB1 | T59S E75D G104E M171V D172E M179I R197K R214K I372M N375S R430K A587P S694N G738E |
| PB1-F2 | G4E T7I Q8P S12L T18I R20K R21K E22G E31G G40D Y42C R44M T46M N47S R48Q A49V R53K Q54R I55T C57S W58L S66N T68I G70E E73K L75R S82L S84N N90S |
| PA | K113R L219I S222N Y241C K254N V261L S277P F354I M441V K497R S558L |
| PA-X | K113R S193N S219F A222T T241A K252R |
| HA | N12S L120M L131Q T156A T211I V226A I526V |
| NP | V105M S450N S482N |
| NA | V67I N71S V206L L269M V321I S339P M412L |
| MP | N82S N85S N87T A227T |
| M2 | R57G D84N |
| NS1 | L7S P83S S87P C116S T127N D139N G204R A223E |
| NEP | L7S G67E |

